# High-resolution dissection of concept acquisition in different families of protein language models

**DOI:** 10.64898/2026.07.20.739599

**Authors:** Shawn Whitfield, Tom Marty, Robert M. Vernon, Christopher James Langmead, Dhanya Sridhar, Quentin Fournier

## Abstract

Protein language models have been increasingly successful on tasks ranging from fitness prediction to functional design, yet what biological knowledge they acquire and where it is encoded within their internal representations remain underexplored. Through a high-resolution layer-by-layer interpretability analysis of 8 models from the ESM2 and AMPLIFY families on 22 concepts from human proteome annotations, we found that these models encode concepts of increasing levels of complexity along their depth: basic physicochemical properties and linear motifs are best captured by early-layer embeddings, secondary structure from subsequent layers, and domain-level semantics from middle layers. Principal component projections of these embeddings showed that they separate biologically meaningful protein groupings, and molecular-biology-inspired interventions demonstrated that pLM embeddings can discriminate phosphomimic-active from inactive mutants. Perhaps surprisingly, we observed that pretraining data and compute had a greater impact on the linear emergence of biological concepts than scaling up parameters. By revealing where biological knowledge is captured in pLMs and which choices shape its emergence, our work offers insights to develop more robust, biologically grounded protein language models.

## Introduction

Proteins are molecular machines that carry out a range of cellular functions. They are composed of linear chains of amino acids, whose chemistry and order dictate their two- and three-dimensional structure, which in turn largely determines their function ^4^. Accurately modeling how sequence relates to structure and function is essential to fundamentally understand biology and discover, manipulate, and design proteins for specific purposes, including breaking down plastics ^66^ or binding to new targets ^40^.

Protein language models (pLMs) have emerged as powerful deep learning tools for modeling these relationships ^23,24,28,36,38,50^. By learning to recover masked amino acid identities from large sequence databases, pLMs encode evolutionary statistics relevant to many aspects of protein function ^68^. Their internal representations, referred to as embeddings, are numerical vectors that are predictive of basic amino acid characteristics, including hydrophobicity, polarity, and size ^23,28,50,60^, protein characteristics including secondary structure ^50^, and protein cellular context including localization signals ^31,45,59^ and membrane integration ^23,45^. pLMs embeddings can encode even higher-level information like gene ontology (GO) annotation ^37^ and evolutionary relatedness ^20,50,60^.

Despite the success of pLM representations learned from coevolutionary statistics, they fail on some downstream tasks and regions of the protein space, suggesting they may not fully capture the fundamental processes underlying protein evolution. For instance, structure-based methods built on pLMs, such as ESMFold ^36^, consistently mispredict protein isoforms, labeling them as structures with aggregation-prone residues ^68^. Recent research using adversarial examples based on well-established physical, chemical, and biological principles found that models like AlphaFold3 and RoseTTAFold tend to overfit their training data rather than following expected physical behaviors ^39^.

To develop pLMs that capture the fundamental mechanisms of biology, the community needs to have a deeper understanding of their learning dynamics and how they encode concepts. In particular, it is important to study how models organize biological information across their layers. While prior studies on pLMs identified that, similarly to natural language models, earlier layers captured “less complex” features such as amino acid biochemistry, middle layers encoded secondary structure and domain features, and later layers captured family information ^1^, they were limited in scope and reported conflicting results. Indeed, depending on model and task, they reported peak predictive performance in both early and late layers ^7,21,35^. Furthermore, interpretability studies, which more formally inspected layerwise pLM embeddings, were mostly restricted to one or two models from the ESM2 model family ^1,25,26,48,53^ and did not examine the impact of data distributions, model size, architecture, or compute. As a result, it remains unclear which biological concepts pLMs acquire, where in the network they emerge, and which factors shape their encoding.

Here, we systematically applied interpretability techniques to evaluate the layerwise embeddings from 8 pLMs from the ESM2 and AMPLIFY families, as well as their structure-aligned variants, across 22 partially overlapping biological concepts. Specifically, we curated datasets of annotated proteins from the human proteome, with labels reflecting their biological functions at multiple levels. Following the observation that language models can linearly encode high-level concepts in their representations ^47^, we trained linear classifiers, referred to as probes, to predict these labels from pLM embeddings at all layers, shedding light on the information they encode. We visually inspected the organization of the embeddings by projecting them using principal component analysis (PCA). Finally, drawing inspiration from molecular biology, we conducted sequence-level interventions to evaluate whether pLMs captured biologically meaningful sequence differences. Together, these analyses show that pLMs encode biological concepts of increasing complexity across depth, with predictive performance often peaking by layer 12. We further find that pretraining data and compute shape concept acquisition more than parameter scaling, and that the resulting embeddings can discriminate phosphomimic-active from inactive mutants.

### Background and related work

In an effort to make this article accessible to readers with backgrounds in either biology or machine learning, we first establish a shared vocabulary that contextualizes our findings, and provide some biological background.

### Proteins

Proteins operate in increasing levels of biological complexity (Figure 1). At a primary level, a protein is defined by its linear sequence of amino acid residues, whose physicochemical properties, such as charge and hydrophobicity, form the basis for subsequent folding. Complexity increases at the secondary level, as the chain forms local motifs like alpha helices and beta sheets. These motifs serve as building blocks for the tertiary level, where the protein folds into a specific three-dimensional shape, creating functional domains. Multiple proteins often assemble into higher-order structures at the quaternary level. Beyond static topology, protein function is a dynamic property modulated by cellular context, including localization and molecular interactions, and temporal scales, ranging from microsecond conformational shifts to evolutionary trajectories.

**Figure 1.**
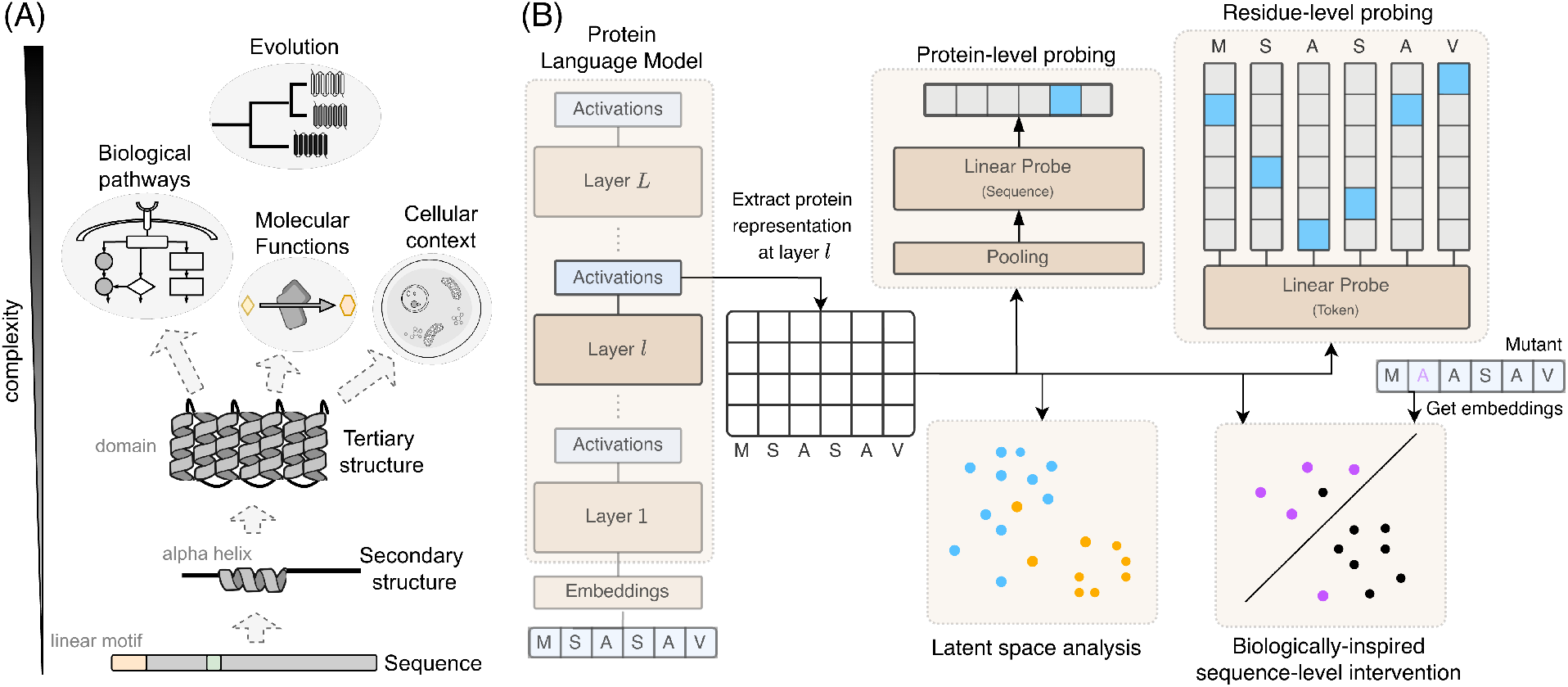
Conceptual overview. A) Proteins work at different levels of biological complexity. Cartoons are modified from NIH BioArt. B) Overview of the interpretability experiments conducted in this study. Proteins are tokenized, and we obtain hidden representations of the protein at each layer (layer *l*). These hidden representations are mean-pooled and used for protein-level linear probing, or used as-is for residue-level linear probing. We also use hidden representations as inputs to PCA analyses to explore the structure and organization of the latent space, and to test whether pLM representations allow discrimination between wild-type and mutant forms of a protein with activity predicted by biological theory.

All this complexity has been captured and distilled by the bioscience community over decades, resulting in organization systems for biological information and protein annotations. This study includes InterPro classification of protein regions ^8^, Gene Ontologies to characterize proteins by Cellular Component (CC), Molecular Function (MF), or Biological Process (BP) ^5,17^, and study-specific information about proteins. Much of this information is collected in the UniProt Knowledgebase ^18^, which provides annotations at various levels of protein biology and rich datasets from which to learn fundamental patterns.

### Protein language models

Protein language models (pLMs) are deep neural networks that learn to represent amino acid residues in a protein as *d*-dimensional vectors, called embeddings. Formally, a pLM with *L* layers takes an amino acid sequence ***x*** = (*x*1, . . . , *x_S_*) of length *S* as input and generates a sequence of residue-level embeddings *f_l_* (***x***) ∈ ℝ*^d^*^×*S*^ at each layer *l*. For left-to-right pLMs, the model is trained to predict the next amino acid *x_s_* by modeling its conditional distribution *P*(*x_s_* |***x***−*s*) as a linear function of the final-layer embedding *f_L_* (***x***−*s*) of the sequence processed so far. For masked pLMs, the network leverages bidirectional context to reconstruct corrupted inputs. Formally, a subset of indices *M* ⊆ {1, . . . , *S*} is selected, and a modified sequence ***̃x*** is generated by replacing the input tokens *x_m_* with a special mask token for all *m* ∈ *M*. The model is then trained to recover the original amino acid identities by modeling the conditional distribution *P*(*x_m_* |***̃x***) as a function of the final-layer embedding *f_L_* (***̃x***)*_m_* at the masked positions. The residue-level embeddings are typically pooled at the final layer, most commonly by averaging across the sequence, to produce a single protein-level embedding ∈ ℝ*^d^*. Intuitively, embeddings should capture a protein’s underlying biology to enable accurate prediction of its unseen parts. The embedding space, also called latent space, is a compressed representation that preserves relevant information and often encodes similar inputs close together.

Protein language models represent proteins differently depending on design choices, including dataset diversity, architecture, and scale, which we investigate by considering pLMs across three families. The popular ESM2 family ranges from 8M to 15B parameters and is trained with the masked language modeling objective (MLM) ^22^ on proteins sampled from UniRef90 based on UniRef50 clusters. The AMPLIFY family ^24^ broadens the protein distribution by training with MLM on UniRef100, Structural Classification of Proteins (SCOP) domains ^2,3^, and Observed Antibody Space (OAS) antibody sequences ^32,46^, while incorporating modern architectural optimizations such as flash attention ^19^ and rotary position embeddings ^57^ for efficiency. Finally, the structure-aligned variants of AMPLIFY models (SaAMPLIFY) use latent-level contrastive learning to align residue representations from pLMs with those from protein graph neural networks ^13^. Despite differences in datasets, architectures, and training methods, all these models aim to capture the multifaceted nature of protein biology.

### Linear probing

A key technique for determining whether an embedding explicitly encodes meaningful properties is linear probing. In this study, we have access to a dataset of different annotations given to proteins. We refer to these annotations as “concepts” to keep with convention in interpretability literature but put simply, a concept here is a classification or categorization one can apply to a protein. Then, we train a linear classifier to predict each annotation given a pLM’s hidden representation at some layer *l*. Formally, given an embedding *f_l_* (***x***) at layer *l*, a probe models the probability of an annotation *c* as *P*(*c*| *f_l_* (***x***)) ∝ exp ***w***⊤ *f_l_* (***x***) , where ***w*** is a learnable weight vector fit on ground-truth annotations. The accuracy of the probe on held-out labeled samples tells us whether the concept is linearly separable given the information the hidden representation encodes. Probes tell us what concept predictions (i.e., categorizations) a pLM can make at each layer based on its hidden representation but do not tell us if the pLM encodes and uses some concept *c* to make its final predictions^1^. The latter is a causal statement that requires intervening on the activations of pLM to observe its effects on a pLM’s predictions. In this study, we are interested in assessing models’ encoding behaviors and the suitability of their representations for a range of concepts, rather than asserting that a model definitively encodes or uses a biological concept. In this context, “concept acquisition” means that the pLM layer representations become potentially useful for predicting annotations of interest.

An important yet sometimes overlooked step to ensure that probes detect genuine signals resulting from training rather than artifacts of the network’s architecture or random initialization is to include robust controls. We argue that probe performance should be contextualized using three kinds of controls, directed against either *input sequences*, *model weights*, or *pLM embeddings* (Table 3). First, *shuffling the input sequence* maintains amino acid composition, removing amino acid ordering and the contribution of positional syntax. Second, *resetting model weights* tests the benefit of learning against random initializations to verify that the pLM learns biologically meaningful information during pre-training, and that the probe is properly trained to detect signals from embeddings. Third, randomly sampling, normalizing, or aggregating (e.g., batch-mean) pLM embeddings tests that associations between proteins and annotations are specific to individual proteins, not to biologically-uninformative characteristics of the latent space or the training process. This last set of controls is motivated by findings in the single-cell perturbation transcriptomics community that a mean baseline often outperforms more complex deep learning approaches ^30,65^. We systematically included these controls.

**Table 1.** Dataset metadata for datasets used in this study. Abbreviations: MF: Molecular Function; BP: Biological Process; CC: Cellular Component; ssp: secondary structure prediction; # sampl.: number of samples; # feat: number of features; avg # ann: average number of annotations (— indicates not applicable). Group min, max indicates the smallest and largest number of proteins sharing the same annotation, while avg grp size indicates the mean number of proteins having the same annotation for the dataset. Norm higher is Yes / No whether a probe trained on normalized embeddings qualitatively showed higher performance than a probe trained on “standard” pLM embeddings, for this dataset (see e.g., Figure 9).

| source | dataset | task level | task type | # sampl. | # feat | avg # ann | group min,max | avg grp size | split train/val/test | norm higher |
| --- | --- | --- | --- | --- | --- | --- | --- | --- | --- | --- |
| uniprot | lipidation | residue | multiclass | 978 | 25 | — | — | — | 0.67/0.13/0.20 | Y |
| uniprot | topology | residue | multiclass | 4777 | 17 | — | — | — | 0.71/0.16/0.14 | Y |
| uniprot | peptide | residue | multiclass | 4977 | 4 | — | — | — | 0.70/0.14/0.16 | Y |
| uniprot | functional sites | residue | multiclass | 6893 | 6 | — | — | — | 0.69/0.16/0.15 | Y |
| uniprot | phosphorylation | residue | multiclass | 8648 | 6 | — | — | — | 0.70/0.15/0.15 | N |
| uniprot | secondary structure | residue | multiclass | 8986 | 4 | — | — | — | 0.70/0.15/0.15 | Y |
| biomap | ssp q3 | residue | multiclass | 11479 | 3 | — | — | — | 0.85/0.09/0.06 | Y |
| biomap | ssp q8 | residue | multiclass | 11479 | 8 | — | — | — | 0.85/0.09/0.06 | Y |
| uniprot | post translational modification | residue | multiclass | 15349 | 12 | — | — | — | 0.70/0.16/0.15 | Y |
| interpro | binding site | protein | binary | 640 | 39 | 1.00 ± 0.06 | [2,298] | 16.46 | 0.70/0.14/0.17 | N |
| interpro | active site | protein | binary | 718 | 58 | 1.19 ± 0.43 | [2,255] | 14.69 | 0.67/0.16/0.17 | N |
| interpro | repeat | protein | binary | 1189 | 65 | 1.28 ± 0.50 | [2,209] | 23.37 | 0.69/0.16/0.15 | N |
| interpro | conserved site | protein | binary | 2948 | 334 | 1.07 ± 0.29 | [2,185] | 9.49 | 0.70/0.15/0.15 | N |
| interpro | family | protein | binary | 3601 | 239 | 1.25 ± 0.53 | [2,716] | 18.89 | 0.58/0.19/0.23 | N |
| biomap | metal ion binding | protein | binary | 7242 | 2 | — | — | — | 0.74/0.08/0.18 | N |
| interpro | domain | protein | binary | 8003 | 364 | 1.64 ± 1.06 | [2,731] | 36.06 | 0.66/0.17/0.17 | N |
| interpro | homologous superfamily | protein | binary | 8198 | 100 | 1.44 ± 0.67 | [13,844] | 117.81 | 0.67/0.17/0.16 | N |
| uniprot | GO MF | protein | binary | 11504 | 123 | 2.43 ± 1.85 | [38,1843] | 227.66 | 0.70/0.16/0.15 | N |
| uniprot | GO BP | protein | binary | 12255 | 304 | 3.15 ± 3.22 | [29,1305] | 127.12 | 0.70/0.15/0.14 | N |
| uniprot | GO CC | protein | binary | 16443 | 164 | 3.64 ± 2.83 | [30,4990] | 365.16 | 0.69/0.15/0.16 | N |
| biomap | localization prediction | protein | multiclass | 7530 | 10 | — | — | — | 0.71/0.08/0.22 | N |
| this study | prot param | protein | regression | 18126 | 31 | — | — | — | 0.70/0.15/0.14 | N |

**Table 2.** Annotation classes to predict for each residue-level dataset.

| Dataset | Non-null classes |
| --- | --- |
| Secondary structure | HELIX, STRAND, TURN |
| Peptide | PROPEP, SIGNAL, TRANSIT |
| Functional sites | ACT_SITE, ACT_SITE-BINDING, BINDING, BINDING-DNA_BIND, DNA_BIND |
| Phosphorylation | 5-glutamyl_glycerylphosphorylethanolamine, Phosphohistidine, Phosphoserine, Phosphothreonine, Phosphotyrosine |
| Topology | Cytoplasmic, Exoplasmic_loop, Extracellular, Intragranular, Lumenal, Lumenal_melanosome, Lumenal_vesicle, Mitochondrial_intermembrane, Mitochondrial_matrix, Mother_cell_cytoplasmic, Nuclear, Perinuclear_space, Peroxisomal, Peroxisomal_matrix, Vacuolar, Vesicular |
| Post-translational modification | CARBOHYD, CARBOHYD-DISULFID, CARBOHYD-DISULFID-MOD_RES, CARBOHYD-LIPID, CARBOHYD-MOD_RES, DISULFID, DISULFID-LIPID, DISULFID-MOD_RES, LIPID, LIPID-MOD_RES, MOD_RES |
| Lipidation | Cholesterol_glycine_ester, GPI-anchor_amidated_alanine, GPI-anchor_amidated_asparagine, GPI-anchor_amidated_aspartate, GPI-anchor_amidated_cysteine, GPI-anchor_amidated_glycine, GPI-anchor_amidated_serine, LIPID, N-myristoyl_glycine, N-palmitoyl_cysteine, N-palmitoyl_glycine, N6-decanoyllysine, N6-myristoyl_lysine, N6-palmitoyl_lysine, O-octanoyl_serine, O-palmitoleoyl_serine, O-palmitoyl_serine, ω-hydroxyceramide_glutamate_ester, Phosphatidylethanolamine_amidated_glycine, Phosphatidylserine_amidated_glycine, S-farnesyl_cysteine, S-geranylgeranyl_cysteine, S-palmitoyl_cysteine, S-stearoyl_cysteine |

**Table 3.** Explanation of different controls used in linear probing experiments.

| Intervention level | Control | Intervention | Tests |
| --- | --- | --- | --- |
| Input protein sequence | Scrambled sequence | Scramble sequence beyond the first residue | That pLMs are linking sequence to the task |
| Model weights (pLM) | Un-trained pLM | For each module in pLM, reset parameters to PyTorch defaults | That a pLM encodes biologically-meaningful information and a probe isn't just picking up noise |
| Model weights (probe) | Un-trained probe | Reset probe weights and biases to PyTorch defaults | That probes are properly trained to detect signals from pLM embeddings and performance isn't just due to random initializations |
| pLM embeddings | Random baseline | Sample from a Gaussian distribution with mean and std given by all embeddings in the batch | That it's the embeddings that give performance, not just the same distribution of values as the embeddings |
| pLM embeddings | Mean embeddings | Set embeddings to the mean of all embeddings in the batch | That performance depends on individual proteins rather than an aggregate representation of a protein |
| pLM embeddings | Normalized embeddings | Normalize residue-level embeddings along hidden dimension | That probes are responding to position and magnitude in embeddings |

### pLM interpretability

Given the successes of pLMs and the richness of available biological annotations, a growing body of research has sought to determine how pLMs internally represent these hierarchies. Interpretability studies on pre-trained pLMs examine what the attention mechanism focuses on ^33,42,56,62^, visualize the protein embedding space ^20,44,51,67^, apply mathematical techniques for analyzing latent space geometry ^49,61^ or attempt to identify neurons or circuits linked to a biological concept ^6,43,63^. More recently, studies employ sparse autoencoders (SAEs) ^1,25,26,41,53^, a linear information extraction technique intended to uncompress pLM embeddings to a sparse, more interpretable form. Sometimes, these studies use large language models ^26,52,53^ to assist in interpreting SAE activations. However, the current scope of pLM interpretability remains limited. Most research focuses almost exclusively on the ESM2 family ^1,26,53^, leaving the impact of alternative architectures, dataset distributions, and training procedures largely unexplored. Furthermore, the field typically relies on a handful of repurposed Bio-ML benchmarking datasets that are known to suffer from biases ^9,58,64^, limiting the ability to draw consistent conclusions. Finally, the inconsistent use of robust controls raises concerns about the validity of reported observations.

To address these limitations, we systematically evaluated 8 pLMs spanning three families and two orders of magnitude in parameter size, along with intermediate checkpoints. Instead of using legacy benchmarks, we curated datasets for 22 biological concepts of various levels, including InterPro families and GO annotations, to rigorously investigate the concepts learned by pLMs. We define the association between pLM embeddings at a given layer and protein annotations for a specific concept as “concept acquisition” and intentionally avoid suggesting that pLMs “understand” biology. Similarly, “learning” does not imply an underlying intelligence. By mapping these concepts to specific layers, we show that pLMs encode biological information at increasing levels of complexity along their depth, in a manner that is consistent across model families but sensitive to some design choices.

## Results

### Many biology annotations are linearly encoded in pLM embeddings

To evaluate the extent to which pLM embeddings encode concepts linearly, we systematically performed linear probing on carefully curated datasets comprising subsets of the human proteome with annotations across all complexity levels. These annotations included biochemical properties based solely on sequence composition (ProtParam, see Methods), residue-wise secondary structure, domain and family memberships, and Gene Ontology annotations (see Table 1). For each protein, we obtained hidden representations from each layer of every model and evaluated the ability of trained probes to predict these annotations. We found that probes could uncover relationships between representations and annotations to some extent for all concepts (Figure 2), although performance was lower for residue-level and for complex, multifactor protein-level tasks such as GO Biological Process prediction. Notably, when either the pLM or the linear probe was replaced with an untrained version, performance dropped to random, indicating that pre-training pLMs encodes these annotated properties in a linearly decodable manner (Figures 9 to 16). Scrambling input sequences affected all predictions except for ProtParam, as expected, since sequence order determines structure and function, whereas ProtParam depends only on amino acid composition. These extensive controls, which exceed those used in most linear probing studies, increased confidence in our findings.

**Figure 2.**
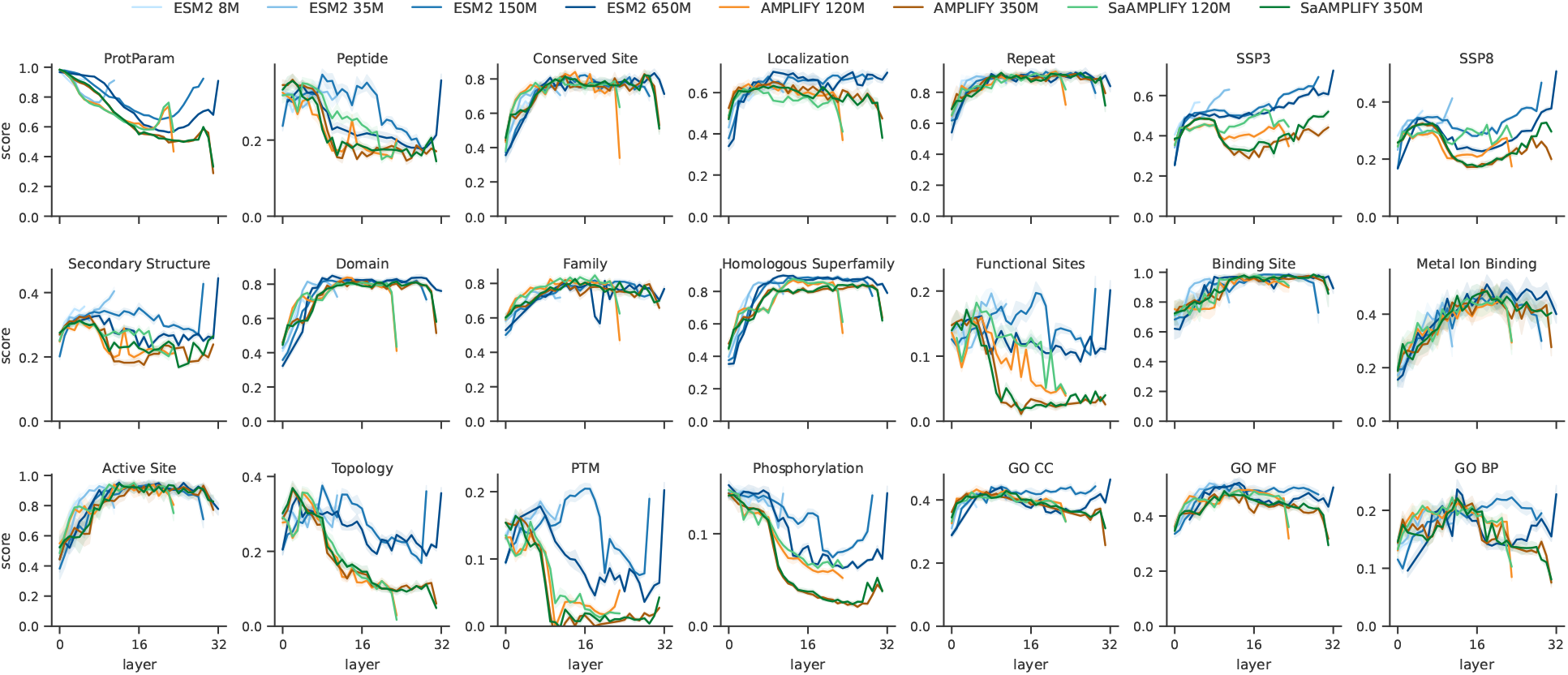
Linear probes trained on pLM embeddings show layer-wise changes in ability to predict annotations covering a range of biological concepts. Score is cosine similarity for ProtParam, otherwise Matthews Correlation Coefficient. Shaded regions show standard deviation from repeated random subsampling 50% of the test set, n=7.

**Figure 3.**
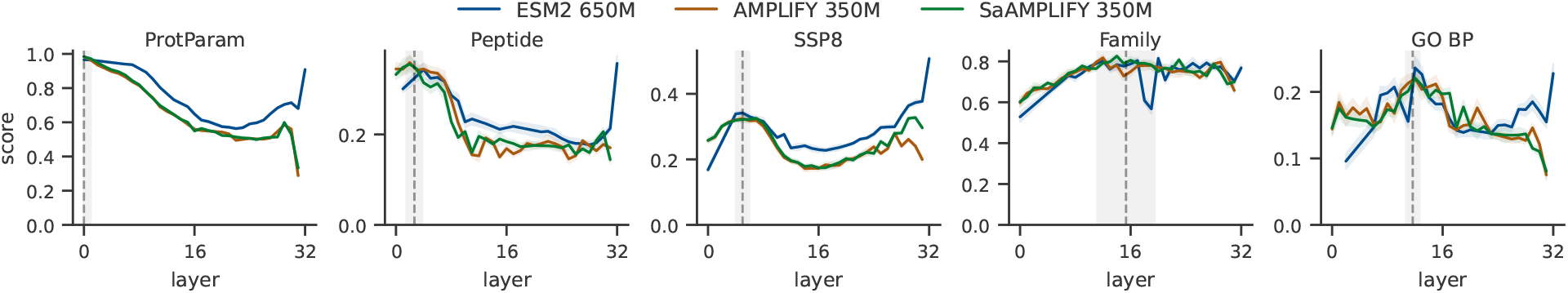
Models share common patterns of concept acquisition, with only minor variations with scale or model type. Horizontal dashed line indicates the average best layer among models shown, and grey shaded region shows the standard deviation of best layers. Score is cosine similarity for ProtParam, otherwise Matthews Correlation Coefficient. Peptide and SSP8 (8-class secondary structure prediction) are residue-level tasks, others are protein-level. Shaded regions along the performance curves are the standard deviation resulting from repeated random subsampling 50% of the test set, n=7.

**Figure 4.**
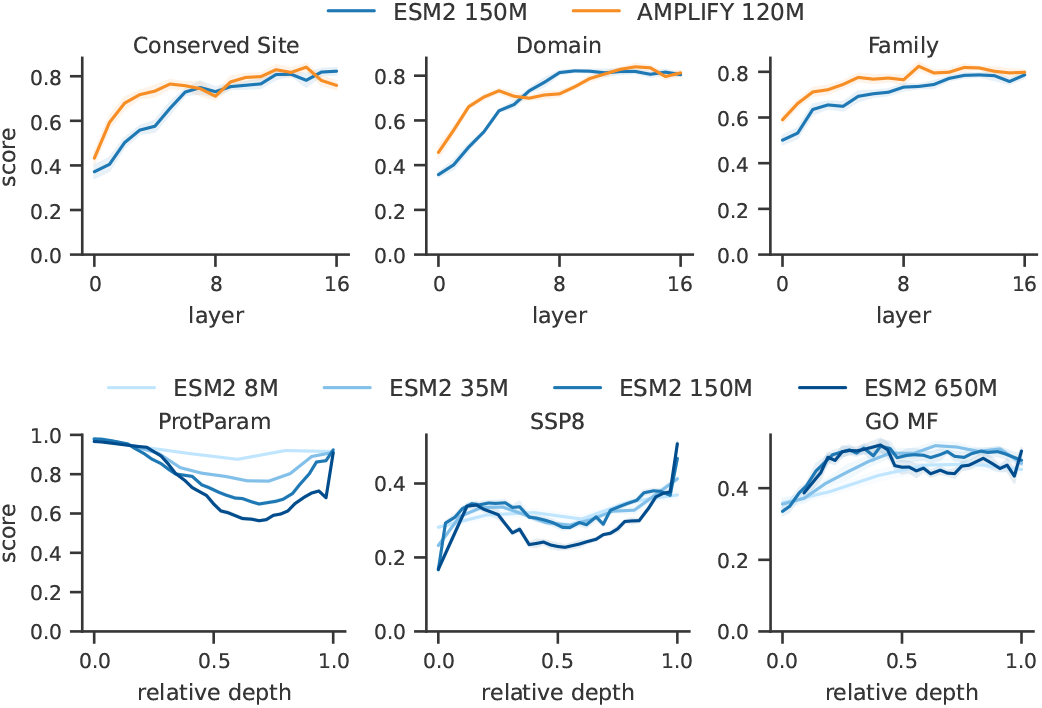
Model family or scale may affect the dynamics of acquiring certain concepts. Top: AMPLIFY models show higher linear probe performance in early layers than ESM2 models of equivalent size. Bottom: Larger ESM2 models perform worse than smaller ones at predicting conceptually simpler protein characteristics (calculated properties using BioPython: ProtParam; 8-class residue-wise secondary structure prediction, SSP8) in later layers, particularly at matched relative depth. This is most apparent when visualizing linear probe performance by relative depth (bottom panels). Shaded regions show standard deviation from repeated random subsampling 50% of the test set, n=7.

**Figure 5.**
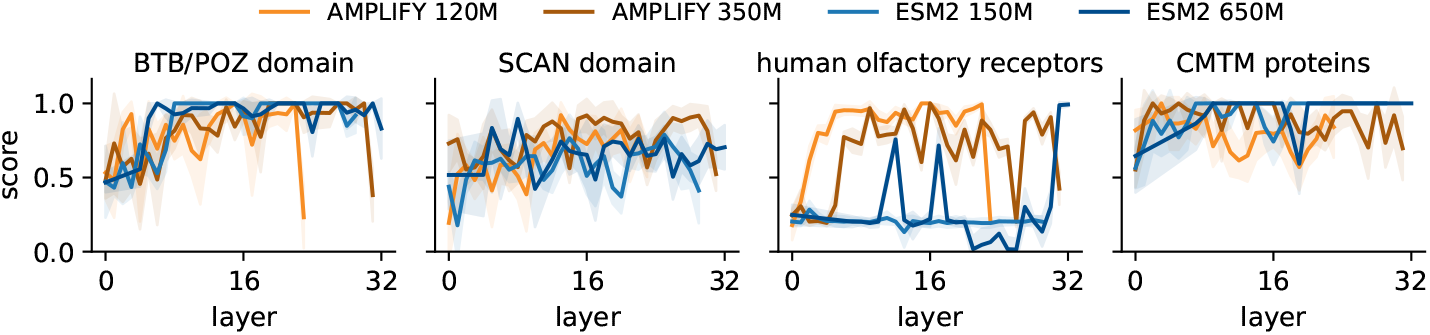
Some specific protein annotations may be more linearly extractable from AMPLIFY or ESM2 embeddings, depending on the layer. Shown here are InterPro annotations: BTB/POZ domain (IPR000210), SCAN domain (IPR003309), human olfactory receptors (IPR050427) and marvel domain-containing and chemokine-like factor (CMTM) proteins (IPR050578). Score is Matthews Correlation Coefficient. Shaded regions along the performance curves are the standard deviation resulting from repeated random subsampling 50% of the test set, n=7.

**Figure 6.**
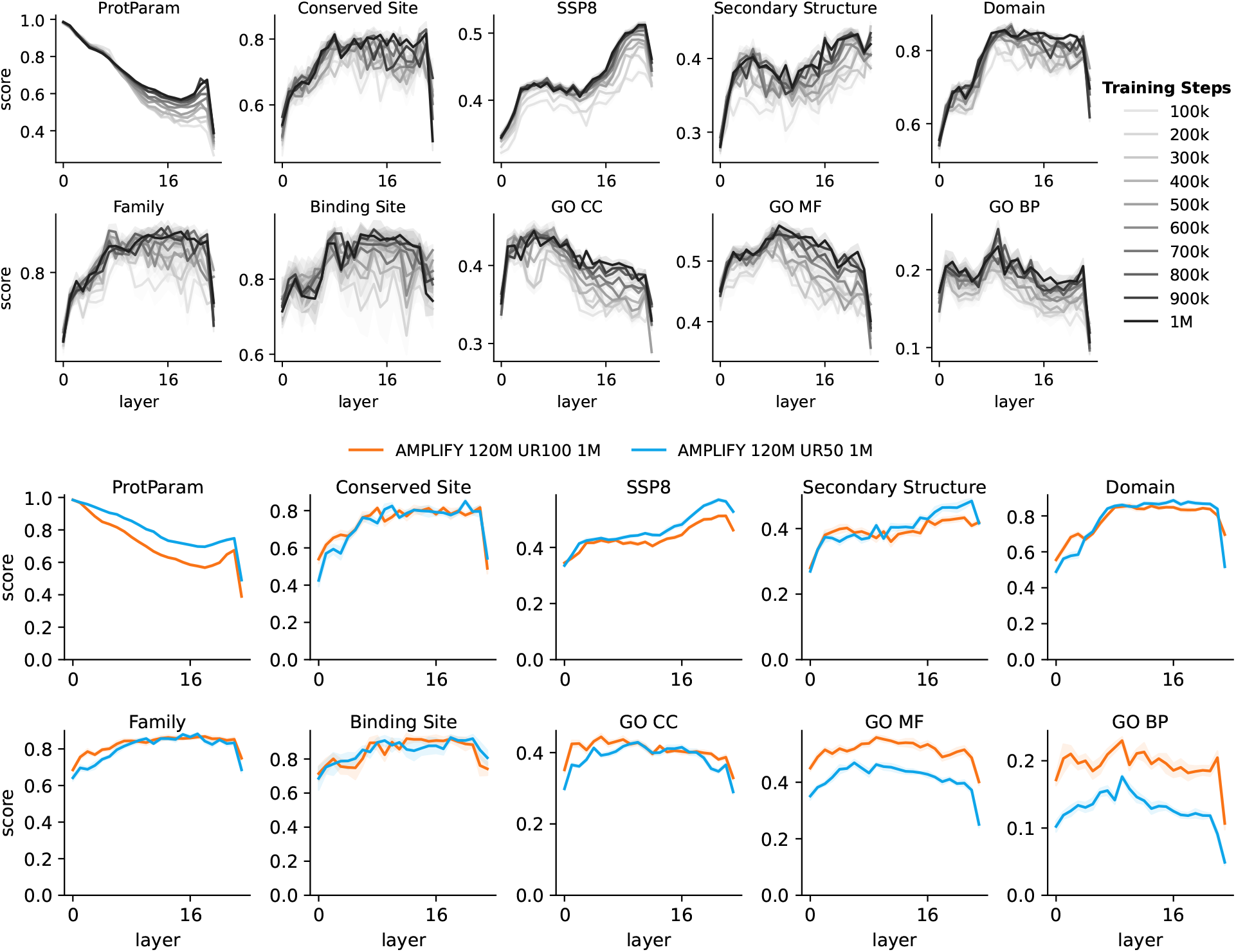
Effect of number of gradient steps on pLM representation performance for different datasets. Datasets were filtered to proteins ≤ 512 amino acids long due to AMPLIFY pretraining strategy. Shaded zones are standard deviation of 50% subsamplings of the test set, n=7. Top: Performance of an AMPLIFY 120M model every 100,000 steps during training. Bottom: Comparison between equivalent layers and checkpoints for AMPLIFY models during training on either UniRef100 (orange) or UniRef50 (blue) (u50).

**Figure 7.**
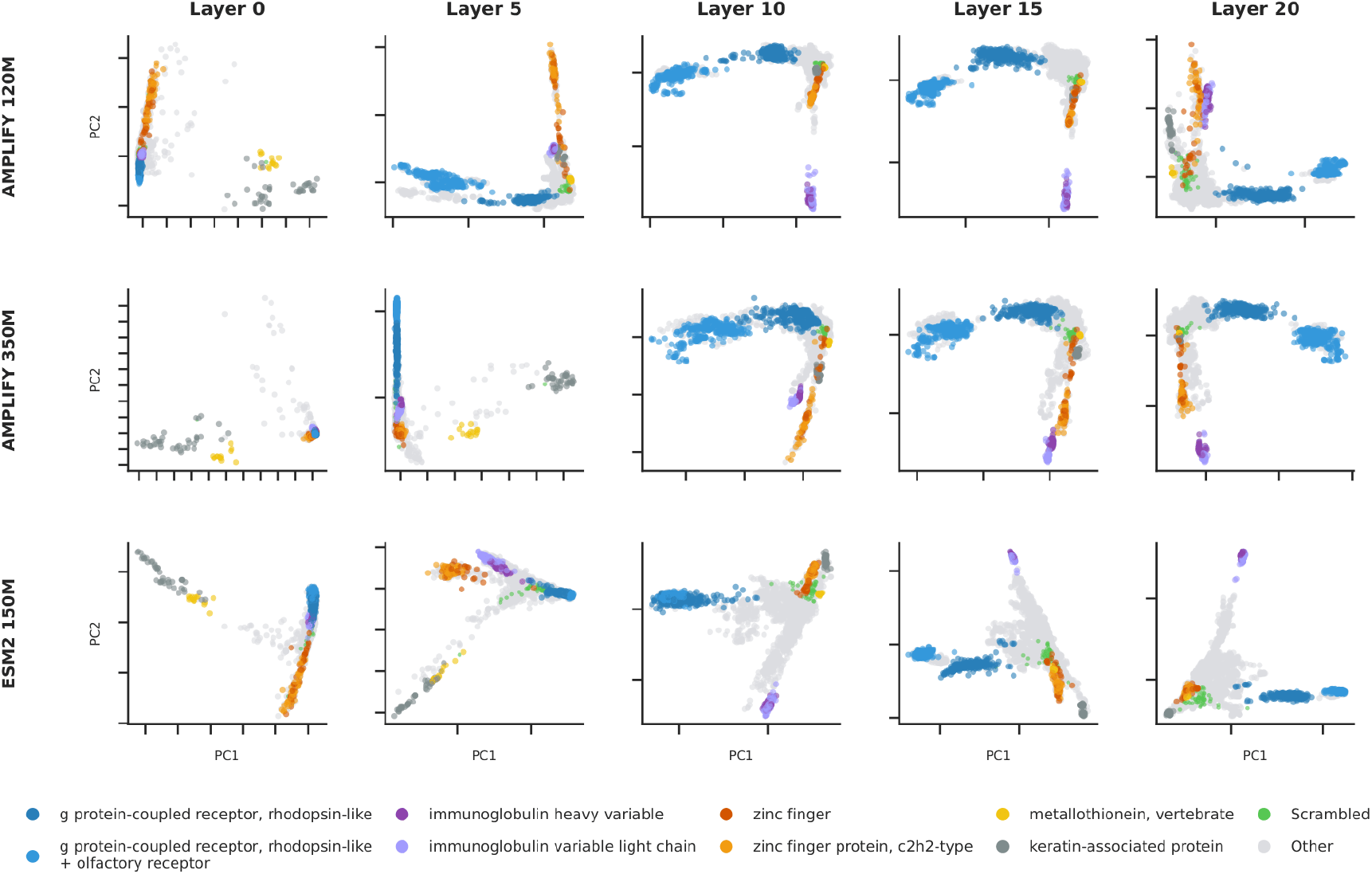
PCA inspection of layerwise embeddings labeled with InterPro Family annotations shows similarities and differences in organization of groupings. Axis ticks denote 20-unit increments.

**Figure 8.**
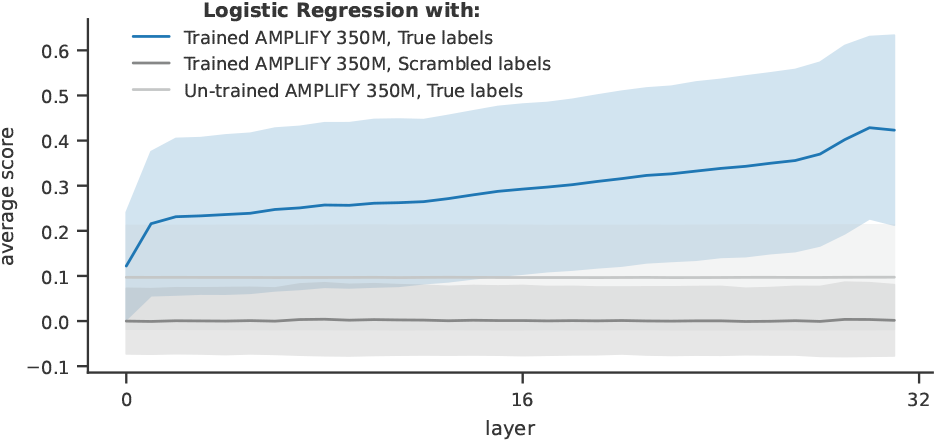
Molecular-biology-inspired manipulations of protein sequences suggest that we can use embeddings to distinguish between theoretically active and inactive forms of a protein. For each randomly-sampled protein (n=1000 samplings), each serine (Ser), threonine (Thr), or tyrosine (Tyr) residue was mutated to alanine (Ala), glycine (Gly), or glutamate (Glu) and a logistic regression classifier trained on a stratified 70:30 split of AMPLIFY 350M embeddings to predict whether a mutant is anticipated to be active (wild-type or Glu at UniProt-annotated phosphosites), inactive (Gly or Ala at annotated phosphosites), or unknown effect (mutations at Ser/Thr/Tyr other than those annotated in UniProt). Average score is the average Matthews Correlation Coefficient among all proteins. Logistic regressors were trained to either predict true labels or scrambled labels from trained AMPLIFY 350M or an un-trained version whose weights and biases were reset to PyTorch defaults. Shaded areas are standard deviation.

**Figure 9.**
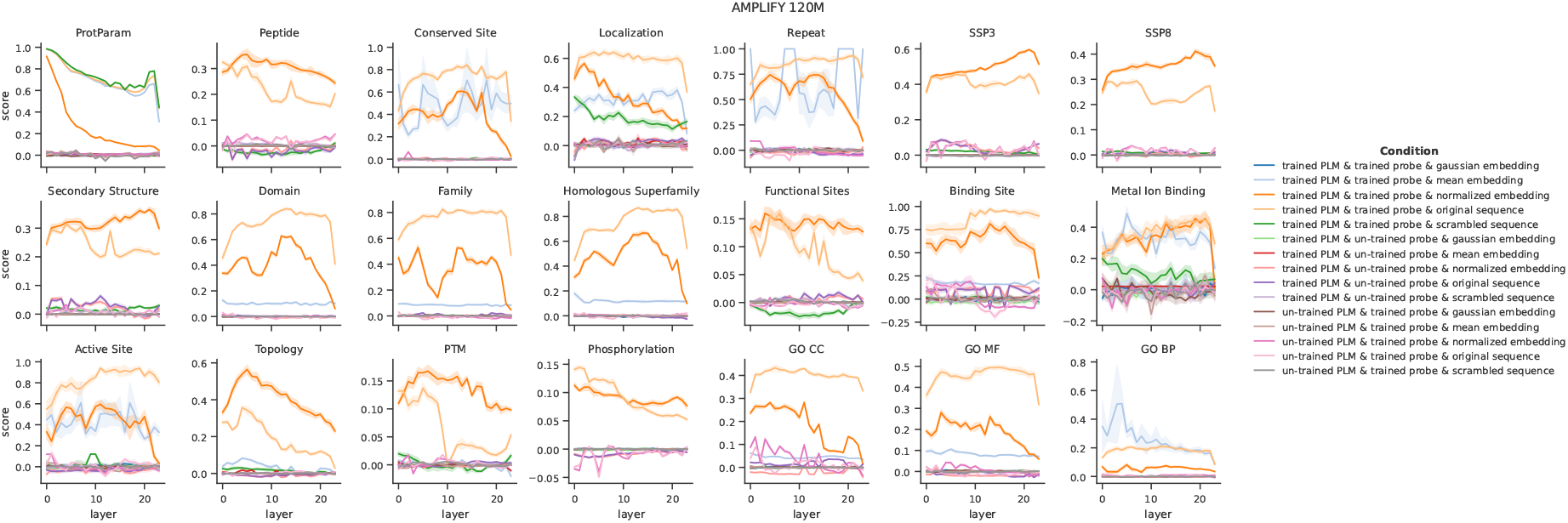
Linear probing controls suggest that probes extract true signal for a range of protein annotations and concepts. Probes were trained either on AMPLIFY 120M embeddings or embeddings from a copy of the model where all weights and biases had been reset to defaults, as a control to verify that pLM embeddings contain meaningful signal that probes can learn (un-trained pLM). Trained probes’ weights and biases were reset to default initializations as a control to verify that probes are capable of learning (un-trained probe). Input protein sequences were either kept as original or scrambled after the first residue to verify that sequence order is important. Random embeddings were drawn from a gaussian with the same distribution as the true pLM embeddings, normalized embeddings were the true embeddings normalized over the pLM hidden dimension, and mean embeddings replaced all embeddings in a batch with the mean vector for that batch. See control details in Table 4.

**Table 4.** How the targets were treated for linear probing controls.

| Control | Task type | Intervention on targets | Reasoning |
| --- | --- | --- | --- |
| Scrambled sequence | Protein-level regression | Calculate ProtParams for new sequence | Easily calculable “ground truth” dependent on new scrambled sequence |
| Scrambled sequence | Protein-level classification | Shuffle target labels for a given protein | Maintain label distribution but unlink sequence from label |
| Scrambled sequence | Residue-level multiclass | Shuffle target labels for a given protein | Maintain label distribution but unlink sequence from label |
| Scrambled sequence | Residue-level binary | Zero-out target labels for a given protein | Scrambling sequence should abolish function |
| Random baseline | All tasks | No change to targets | The input sequence should still map to the same target labels |
| Mean embeddings | Regression | Take the mean for each target property in the batch | A mean representation for the batch should have the mean response for each property in the batch |
| Mean embeddings | Classification | Take the mode for each target label in the batch | A mean representation for the batch should have the most common labels in the batch |
| Normalized embeddings | All tasks | No change to targets | The input sequence should still map to the same target labels |

### Common patterns in concept acquisition among different models

Transformer models are believed to develop lower-level features in earlier layers and encode increasingly complex representations as depth increases. Consistent with this, we found that early-layer embeddings predicted lower-level protein properties such as molecular weight (see Methods and Figure 17) well across various models and scales (see Figures 2 and 3). Predictive performance for annotations associated with short, contiguous protein regions such as repeats, peptide signals, localization, and secondary structure, peaked in subsequent layers, while annotations such as domain, binding site, and protein family peaked several layers later. For most models, we observed a bimodal pattern in secondary-structure prediction: a peak in performance in early layers followed by a decrease/plateau that steadily increased again with depth. Performance on GO annotations generally peaked around the middle third of model layers, if at all. These findings suggest that the models evaluated here, which are likely limited in capacity given the incredible richness of biological information, are forced to make trade-offs rather than systematically build higher-level concepts from lower-level ones. Strikingly, across all models, scales, and datasets, predictive performance on most concepts peaked by layer 12. This observation aligns with a previous interpretability study, which reported a peak in linear probe performance at layer 12 of ESM2 650M across multiple datasets ^1^, from secondary structure prediction and protein localization. This pattern may indicate that, beyond layer 12, models perform complex nonlinear representational adjustments that are not fully detected by our experimental approach.

**Figure 10.**
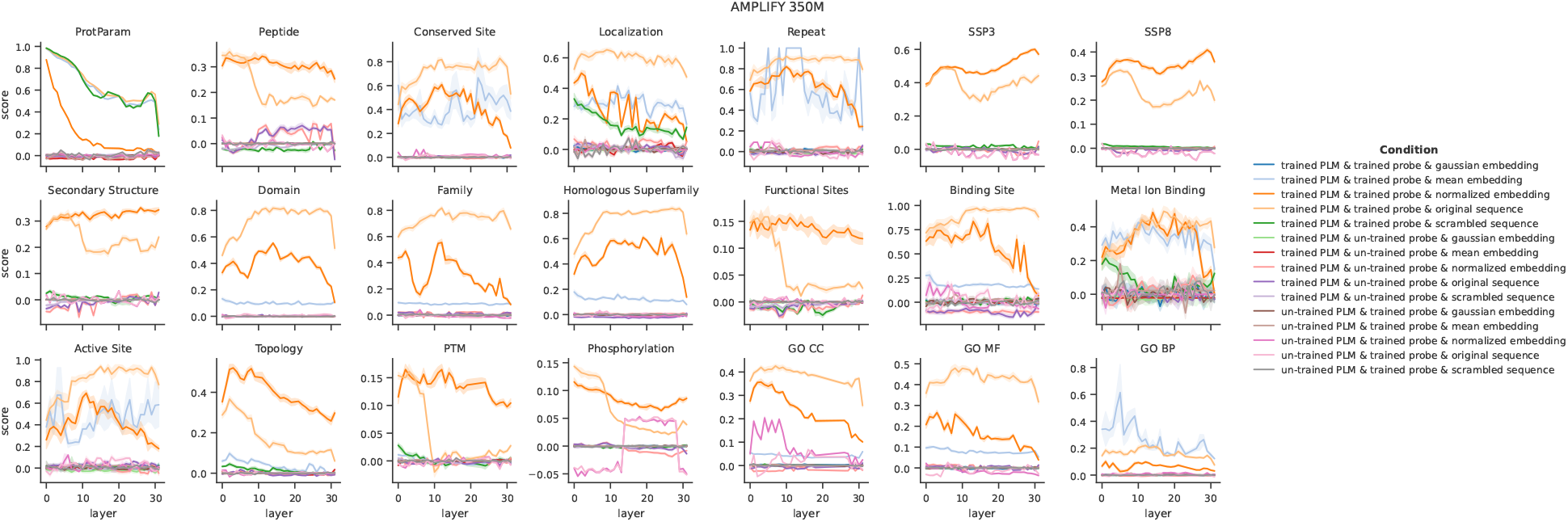
Linear probing controls suggest that probes extract true signal for a range of protein annotations and concepts. Probes were trained either on AMPLIFY 350M embeddings or embeddings from a copy of the model where all weights and biases had been reset to defaults, as a control to verify that pLM embeddings contain meaningful signal that probes can learn (un-trained pLM). Trained probes’ weights and biases were reset to default initializations as a control to verify that probes are capable of learning (un-trained probe). Input protein sequences were either kept as original or scrambled after the first residue to verify that sequence order is important. Random embeddings were drawn from a gaussian with the same distribution as the true pLM embeddings, normalized embeddings were the true embeddings normalized over the pLM hidden dimension, and mean embeddings replaced all embeddings in a batch with the mean vector for that batch. See control details in Table 4.

**Figure 11.**
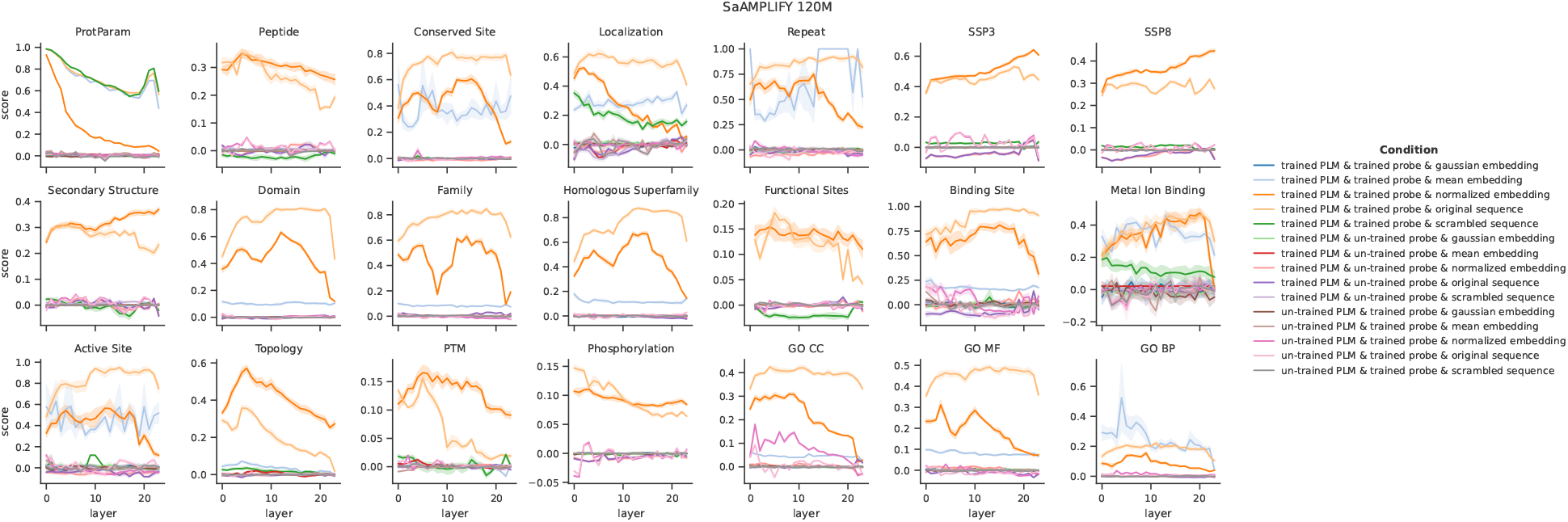
Linear probing controls suggest that probes extract true signal for a range of protein annotations and concepts. Probes were trained either on SaAMPLIFY 120M embeddings or embeddings from a copy of the model where all weights and biases had been reset to defaults, as a control to verify that pLM embeddings contain meaningful signal that probes can learn (un-trained pLM). Trained probes’ weights and biases were reset to default initializations as a control to verify that probes are capable of learning (un-trained probe). Input protein sequences were either kept as original or scrambled after the first residue to verify that sequence order is important. Random embeddings were drawn from a gaussian with the same distribution as the true pLM embeddings, normalized embeddings were the true embeddings normalized over the pLM hidden dimension, and mean embeddings replaced all embeddings in a batch with the mean vector for that batch. See control details in Table 4.

**Figure 12.**
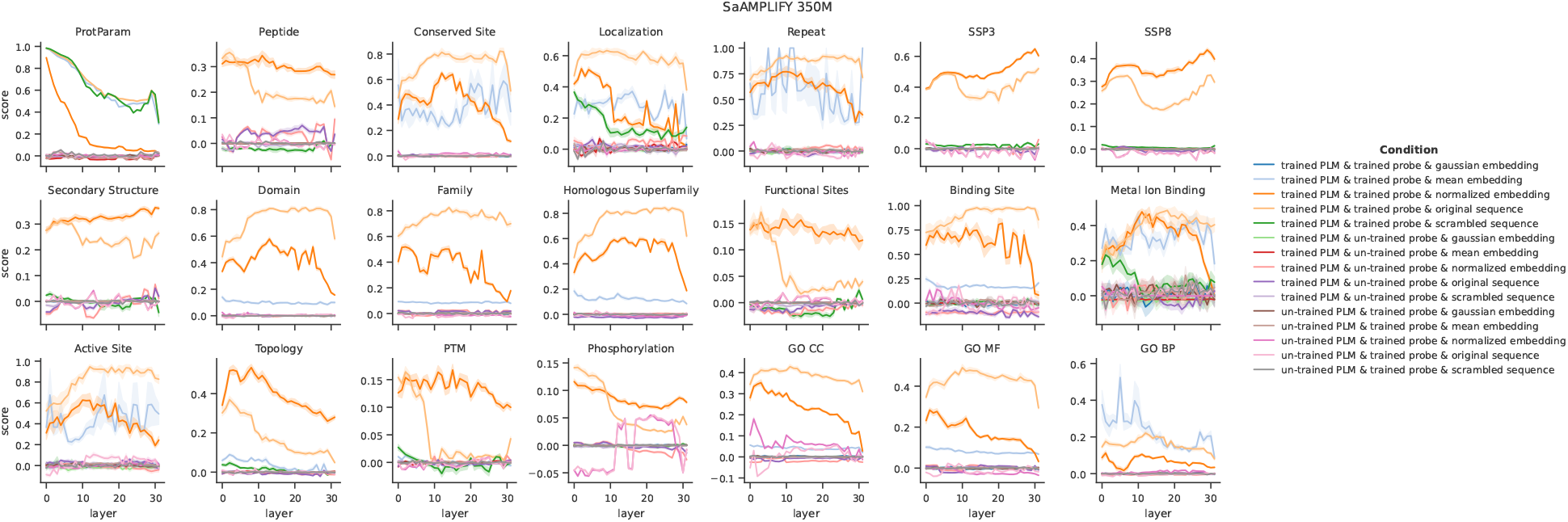
Linear probing controls suggest that probes extract true signal for a range of protein annotations and concepts. Probes were trained either on SaAMPLIFY 350M embeddings or embeddings from a copy of the model where all weights and biases had been reset to defaults, as a control to verify that pLM embeddings contain meaningful signal that probes can learn (un-trained pLM). Trained probes’ weights and biases were reset to default initializations as a control to verify that probes are capable of learning (un-trained probe). Input protein sequences were either kept as original or scrambled after the first residue to verify that sequence order is important. Random embeddings were drawn from a gaussian with the same distribution as the true pLM embeddings, normalized embeddings were the true embeddings normalized over the pLM hidden dimension, and mean embeddings replaced all embeddings in a batch with the mean vector for that batch. See control details in Table 4.

**Figure 13.**
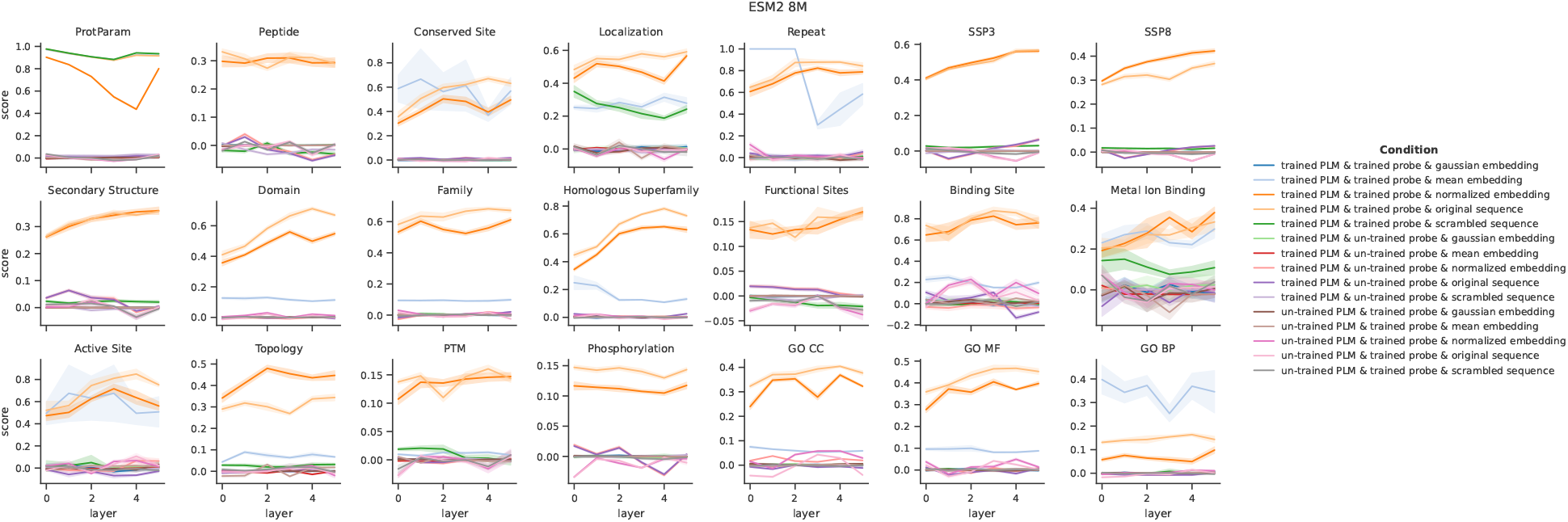
Linear probing controls suggest that probes extract true signal for a range of protein annotations and concepts. Probes were trained either on ESM2 8M embeddings or embeddings from a copy of the model where all weights and biases had been reset to defaults, as a control to verify that pLM embeddings contain meaningful signal that probes can learn (un-trained pLM). Trained probes’ weights and biases were reset to default initializations as a control to verify that probes are capable of learning (un-trained probe). Input protein sequences were either kept as original or scrambled after the first residue to verify that sequence order is important. Random embeddings were drawn from a gaussian with the same distribution as the true pLM embeddings, normalized embeddings were the true embeddings normalized over the pLM hidden dimension, and mean embeddings replaced all embeddings in a batch with the mean vector for that batch. See control details in Table 4.

**Figure 14.**
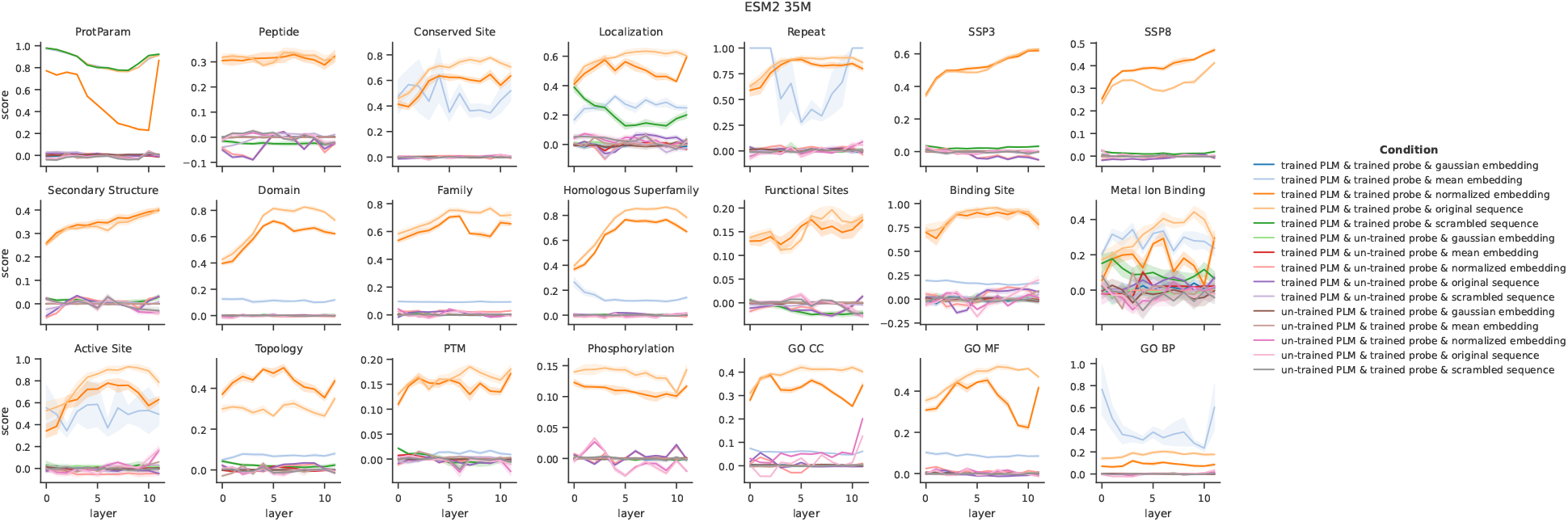
Linear probing controls suggest that probes extract true signal for a range of protein annotations and concepts. Probes were trained either on ESM2 35M embeddings or embeddings from a copy of the model where all weights and biases had been reset to defaults, as a control to verify that pLM embeddings contain meaningful signal that probes can learn (un-trained pLM). Trained probes’ weights and biases were reset to default initializations as a control to verify that probes are capable of learning (un-trained probe). Input protein sequences were either kept as original or scrambled after the first residue to verify that sequence order is important. Random embeddings were drawn from a gaussian with the same distribution as the true pLM embeddings, normalized embeddings were the true embeddings normalized over the pLM hidden dimension, and mean embeddings replaced all embeddings in a batch with the mean vector for that batch. See control details in Table 4.

**Figure 15.**
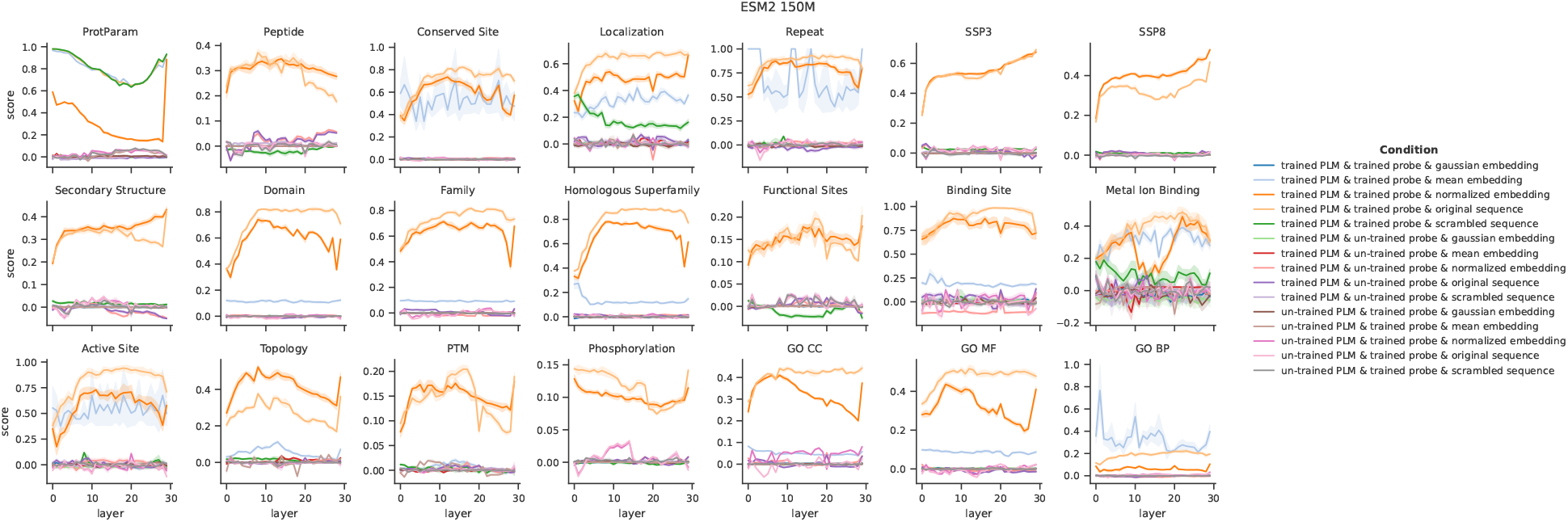
Linear probing controls suggest that probes extract true signal for a range of protein annotations and concepts. Probes were trained either on ESM2 150M embeddings or embeddings from a copy of the model where all weights and biases had been reset to defaults, as a control to verify that pLM embeddings contain meaningful signal that probes can learn (un-trained pLM). Trained probes’ weights and biases were reset to default initializations as a control to verify that probes are capable of learning (un-trained probe). Input protein sequences were either kept as original or scrambled after the first residue to verify that sequence order is important. Random embeddings were drawn from a gaussian with the same distribution as the true pLM embeddings, normalized embeddings were the true embeddings normalized over the pLM hidden dimension, and mean embeddings replaced all embeddings in a batch with the mean vector for that batch. See control details in Table 4.

**Figure 16.**
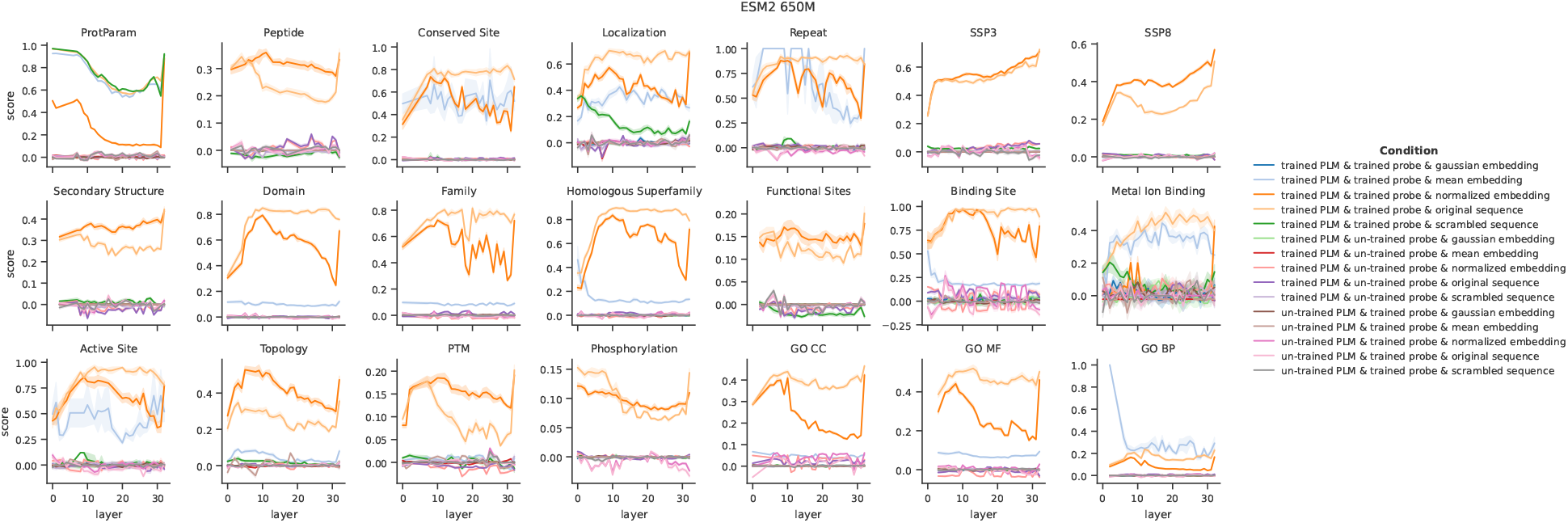
Linear probing controls suggest that probes extract true signal for a range of protein annotations and concepts. Probes were trained either on ESM2 650M embeddings or embeddings from a copy of the model where all weights and biases had been reset to defaults, as a control to verify that pLM embeddings contain meaningful signal that probes can learn (un-trained pLM). Trained probes’ weights and biases were reset to default initializations as a control to verify that probes are capable of learning (un-trained probe). Input protein sequences were either kept as original or scrambled after the first residue to verify that sequence order is important. Random embeddings were drawn from a gaussian with the same distribution as the true pLM embeddings, normalized embeddings were the true embeddings normalized over the pLM hidden dimension, and mean embeddings replaced all embeddings in a batch with the mean vector for that batch. See control details in Table 4.

**Figure 17.**
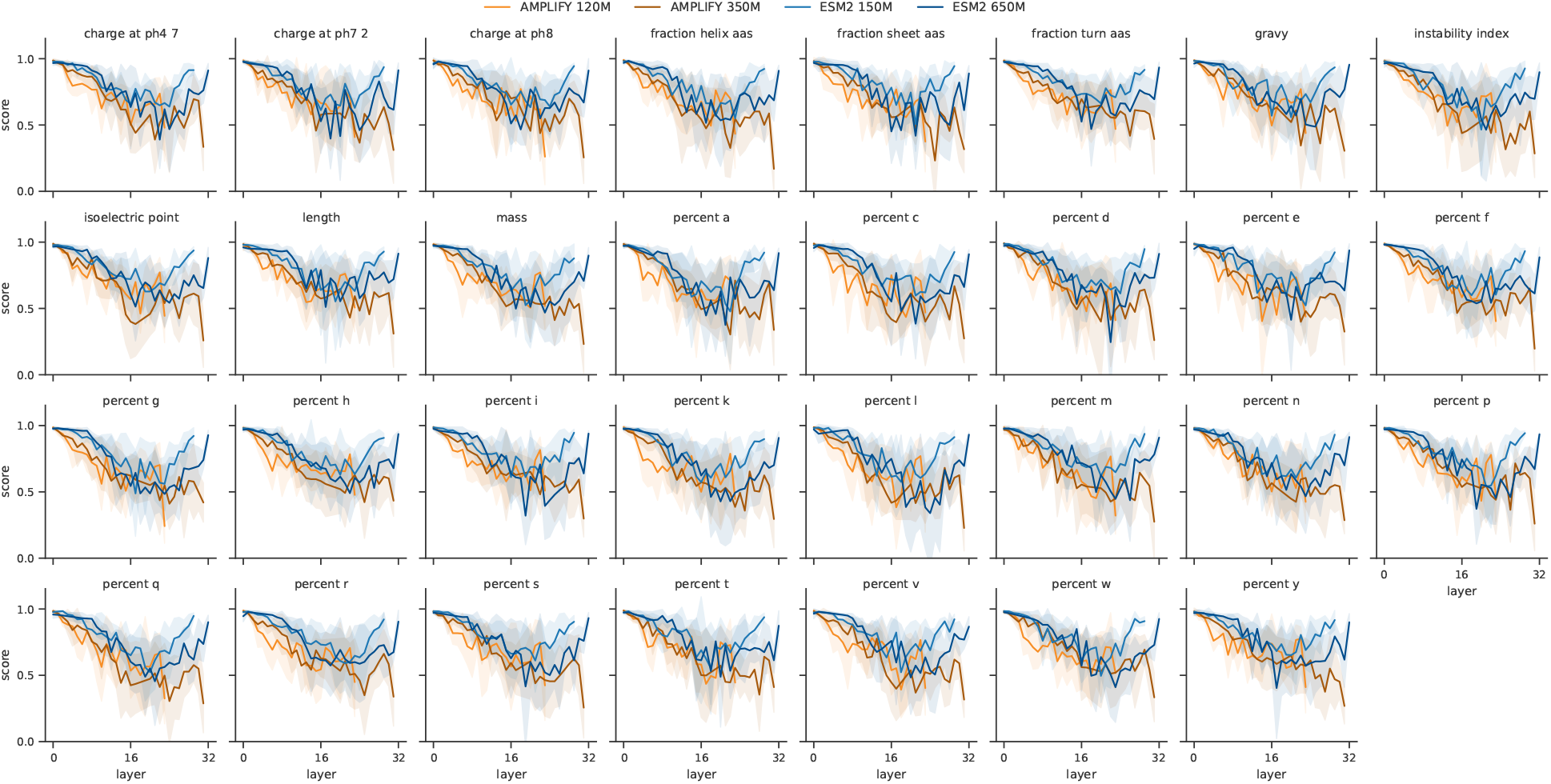
Elementwise results for ProtParam properties, calculated using BioPython’s ProteinAnalysis tool. Shaded regions are standard deviation resulting from repeated random subsampling 50% of the test set, n=7. Score is cosine similarity.

**Figure 18.**
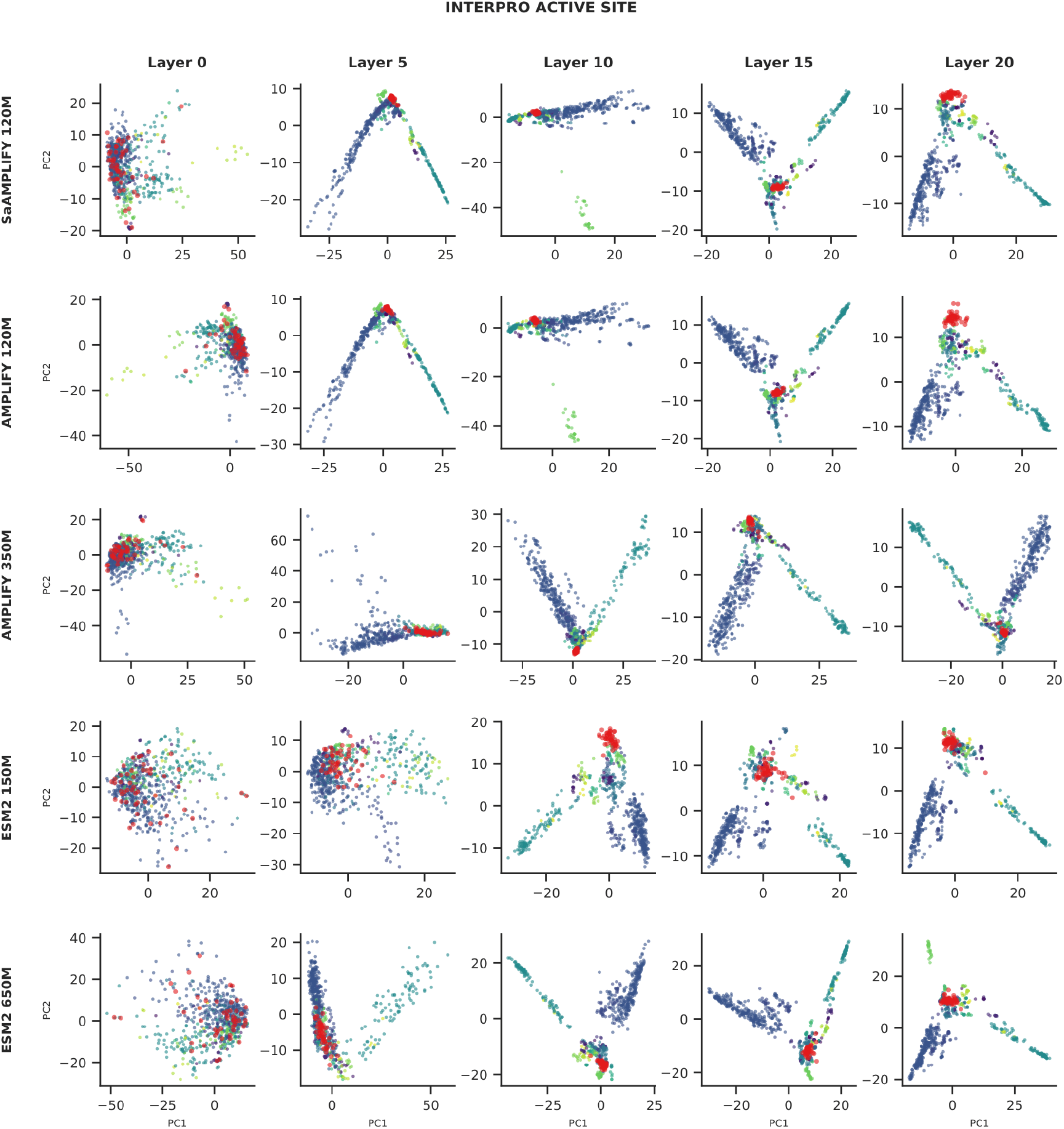
PCA inspection of layerwise embeddings labeled with InterPro Active Site annotations shows similarities and differences in organization of groupings. Axis ticks denote 20-unit increments.

### pLMs acquire some concepts differently based on model family, scale, or training procedure

Although probe performance on datasets such as repeats, metal ion binding, and GO cellular compartments showed limited variation across pLMs, distinct patterns observed in other datasets suggest that models tend to acquire representations of specific concepts preferentially (Figure 4). We highlight these differences below.

### AMPLIFY models acquire certain concepts within fewer layers than ESM2 models, regardless of scale

For several datasets, embeddings from AMPLIFY models consistently showed greater predictive power at earlier layers compared to ESM2 models of similar size (Figure 4, top). This observation suggests that AMPLIFY models may preferentially acquire these concepts due to architectural or training differences. If pLMs encode concepts at increasing levels of complexity along their depth, as we find in this work, a model that learns a given concept earlier should be overall more powerful and potentially useful for more tasks, but more experiments to support this are outside our current scope. Given their architectural similarities, the most plausible explanation is that differences in the training data account for most of the observed differences; however, training both models on identical datasets is beyond the scope of this study.

### Scale is beneficial for the acquisition or maintenance of certain concepts, but not all

Interpretability studies that conducted scale comparisons found that larger ESM2 models captured more interpretable concepts ^53^. While this trend was generally observed in the present analysis, particularly for protein domain annotations, the larger ESM2 model performed worse than the smaller model at later layers, especially for a given relative depth (Figure 4, bottom). These findings suggest that the representation of higher-order and more fundamental processes may be partially mutually exclusive.

### Training cements acquisition of concepts

Following the above findings that model family and scale influence concept acquisition, we further examined the impact of training duration. The AMPLIFY models follow a two-phase training process: an initial phase of 1 million gradient steps with a maximum protein length of 512 residues, during which longer proteins were randomly truncated, followed by a 25 thousand step context extension phase with a maximum protein length of 2048 residues ^24^. Training checkpoints were collected every 100k steps from the initial training phase of a 120M parameter model trained on UniRef100 (Figure 6). Performance improved consistently with additional training steps, with the most substantial gains observed in mid and late layers, while early layers showed minimal variation across checkpoints. These findings suggest that increasing the number of training steps largely contributes to representation improvement rather than enabling the acquisition of entirely new concepts.

### Differences in specific annotations

We evaluated the performance of linear probes on individual protein-level annotations to assess the respective strengths of AMPLIFY and ESM2. We identified instances where linear probes more effectively associated layers of each pLM family with specific annotations (Figure 5). For instance, intermediate layers of AMPLIFY outperformed ESM2 in predicting human olfactory receptors and zinc-finger SCAN domains. These observations support the hypothesis that differences in concept acquisition likely result from the composition of the training datasets. Indeed, the higher proportion of human proteins in UniRef100 (∼178k, 0.0479%), which constitutes most of AMPLIFY’s training set, may provide AMPLIFY with greater exposure to relevant features for these annotations compared to ESM2, which is trained on the less human-rich distribution of UniRef50 (∼12k, 0.0192%).

To assess the impact of training data composition on concept acquisition, we obtained training checkpoints for two small-scale AMPLIFY-style models trained on UniRef50 and UniRef100 (Figure 6, bottom). The model trained on UniRef50 generally performed worse than, or comparably to, the model trained on UniRef100, suggesting that data diversity is key to concept acquisition and further supporting our hypothesis. Performance differences were most evident in GO annotation prediction, indicating that diversity is particularly important for representing higher-level concepts.

### Visualizing how concepts are organized in the latent space

Dimensionality reduction is an effective approach for interpreting the organization of the latent space when labeled data are available. Although UMAP (Uniform Manifold Approximation and Projection) is commonly employed for this purpose, there may be concerns about the validity of conclusions drawn from its results ^11,29^ due to its nonlinear, nondeterministic nature and the sensitivity of the visualization to its hyperparameters. In this study, we instead relied on PCA for visualization because it is faster, reproducible, and more interpretable. Visual inspection of layerwise embeddings annotated by protein family (Figure 7) shows that AMPLIFY 120M and 350M at layers 10 and 15 exhibit similar structures, whereas layers 0 and 5 display distinct organizations. Although ESM2 150M separates many of the same groups as the AMPLIFY models by layer 15, such as immunoglobulin variable chains, olfactory receptors, and G-protein coupled receptors, the structure of their projected embedding spaces appears different. These visualizations support the linear separability of many annotations, as two principal components are often sufficient to distinguish groupings in the embedding space (see Figures 18 to 25 for PCA of other annotations).

### Molecular-biology-inspired interventions suggest pLM embeddings can help distinguish plausible mutants

A common technique used by molecular biologists to investigate protein function is to manipulate, or mutate, a protein sequence to confirm or disprove hypotheses regarding specific activities. To evaluate the sensitivity of pLM embeddings for predicting biologically relevant outcomes, we performed targeted sequence-level interventions. We focused on Phosphorylation, the addition of a negatively charged phosphoryl group to specific residues, which can either activate or deactivate the protein. We chose phosphorylation because there are well-established mutations that a molecular biologist would make: substituting a phosphorylatable serine, threonine, or tyrosine with glutamate often mimics the negative charge and bulk of a phosphoryl group, whereas replacing the residue with alanine or glycine removes the phosphorylatable site without changing protein length. For most other concepts, only interventions that are presumed to inactivate the protein are available.

If embeddings capture information about protein activation, they should allow distinguishing between mutants predicted by biologists as active, inactive, or unknown, depending on the type of mutation and whether the mutated residue is annotated in UniProt as phosphorylated. We tested this hypothesis by training logistic regression classifiers to predict the labels for all possible phosphomutants across 1000 randomly selected proteins. Embeddings from the later layers of AMPLIFY 350M were generally predictive of potential activation status (Figure 8). We observed similar but less striking trends for AMPLIFY 120M, ESM2 150M, and ESM2 650M (see Figures 26 to 28). Shuffling the labels prevented logistic regressors from correctly associating mutants with activation status (highest p-value: layer 0, 6.24e-15; lowest: layer 31, 8.28e-24; using a paired Wilcoxon signed-rank test with Bonferroni post-hoc test to correct for multiple comparisons between layers), indicating that the classifiers learned an association between representations and activation. These findings suggest that, in certain cases, pLM embeddings can be a valuable tool for *in silico* triage of mutants prior to *in vitro* testing.

**Figure 19.**
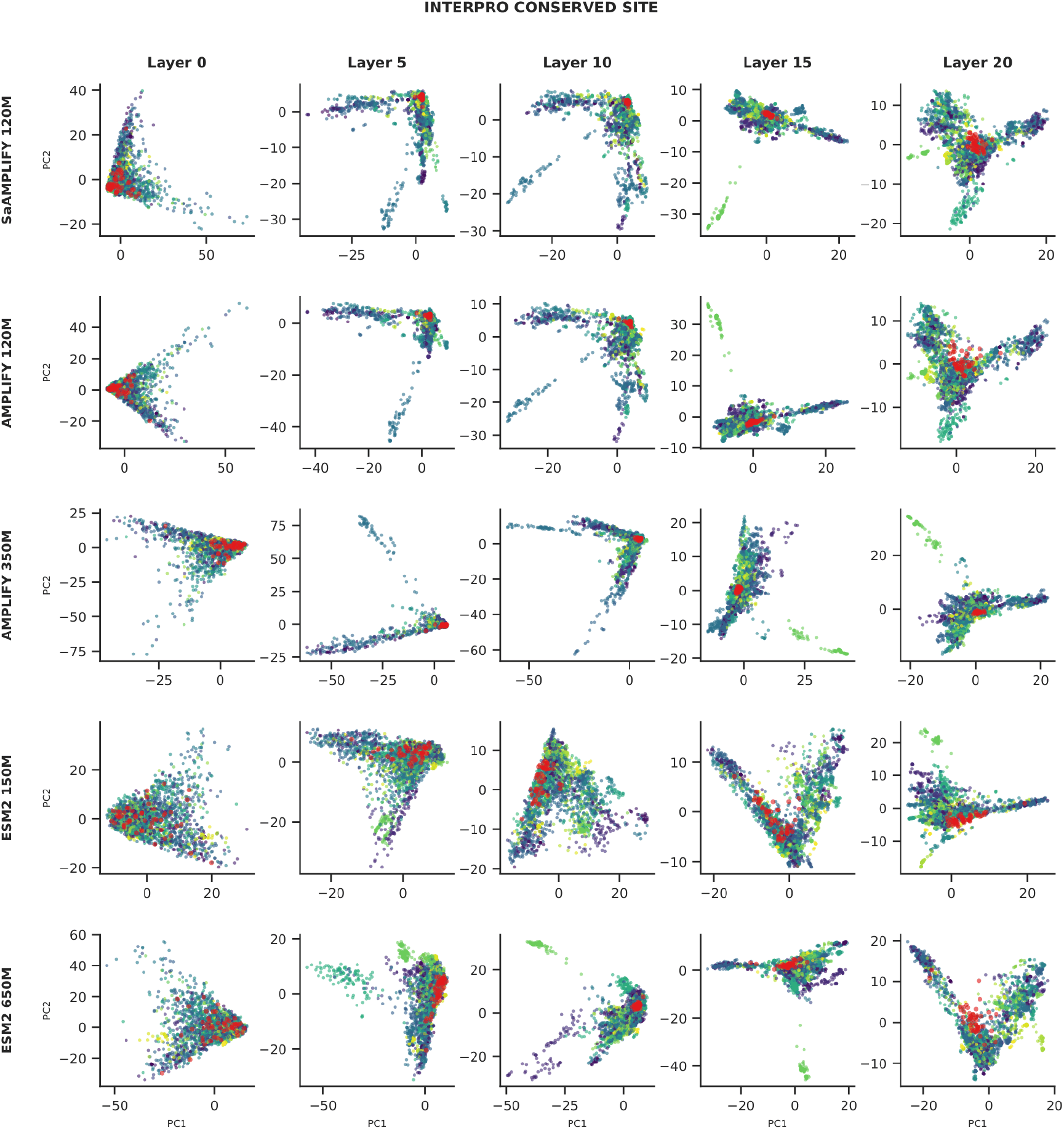
PCA inspection of layerwise embeddings labeled with InterPro Conserved Site annotations shows similarities and differences in organization of groupings. Axis ticks denote 20-unit increments.

**Figure 20.**
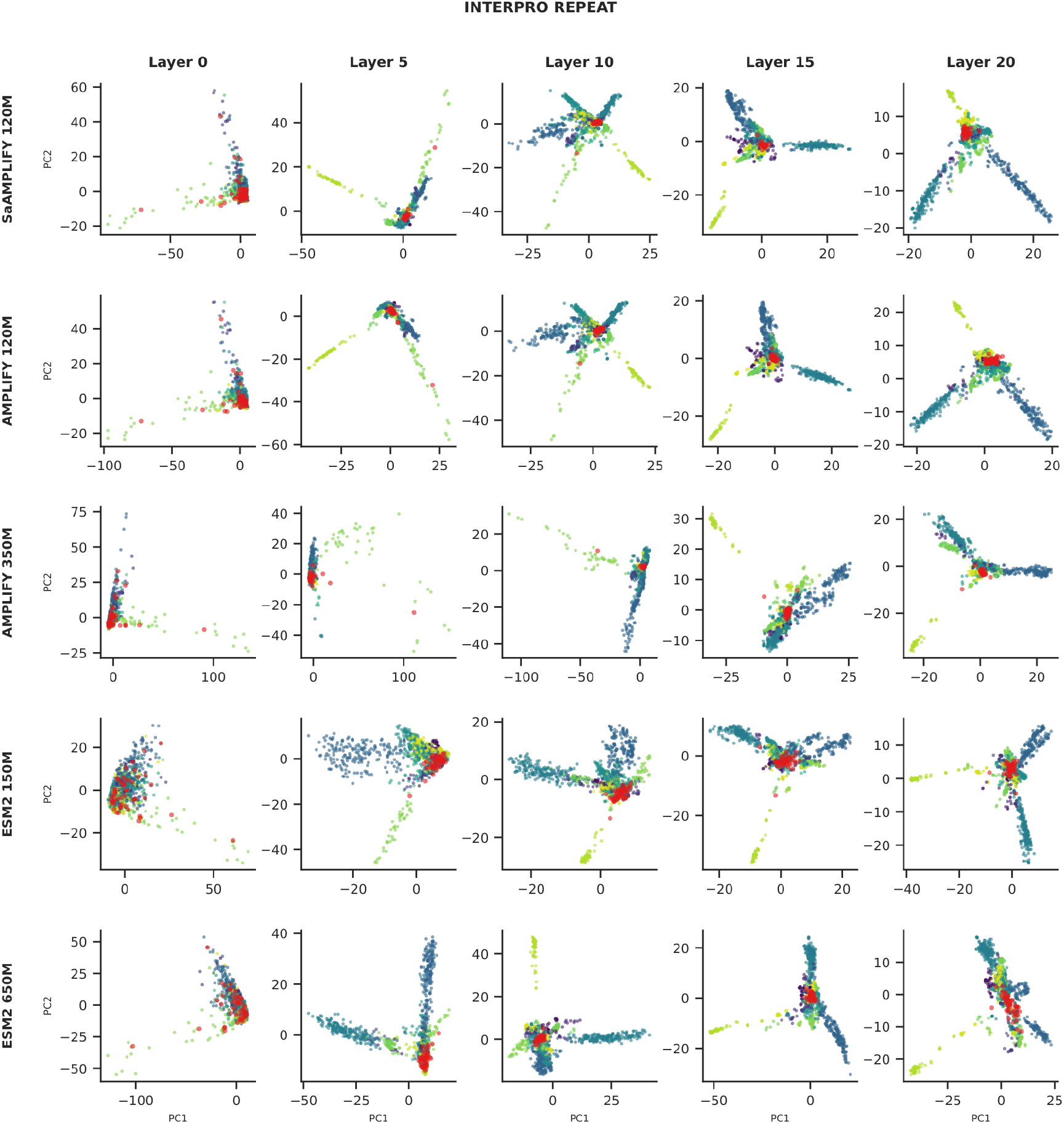
PCA inspection of layerwise embeddings labeled with InterPro Repeat annotations shows similarities and differences in organization of groupings. Axis ticks denote 20-unit increments.

**Figure 21.**
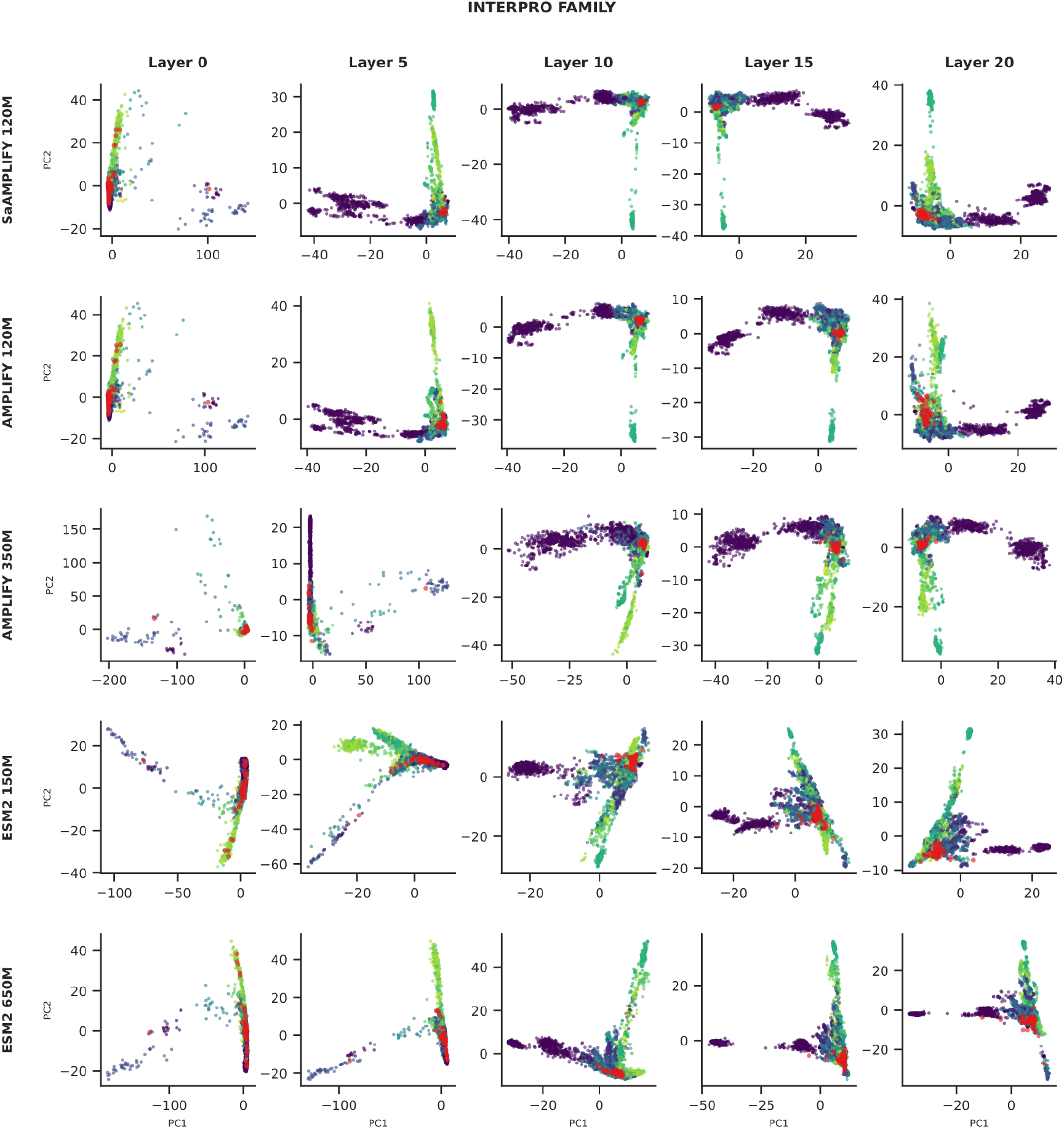
PCA inspection of layerwise embeddings labeled with InterPro Domain annotations shows similarities and differences in organization of groupings. Axis ticks denote 20-unit increments.

**Figure 22.**
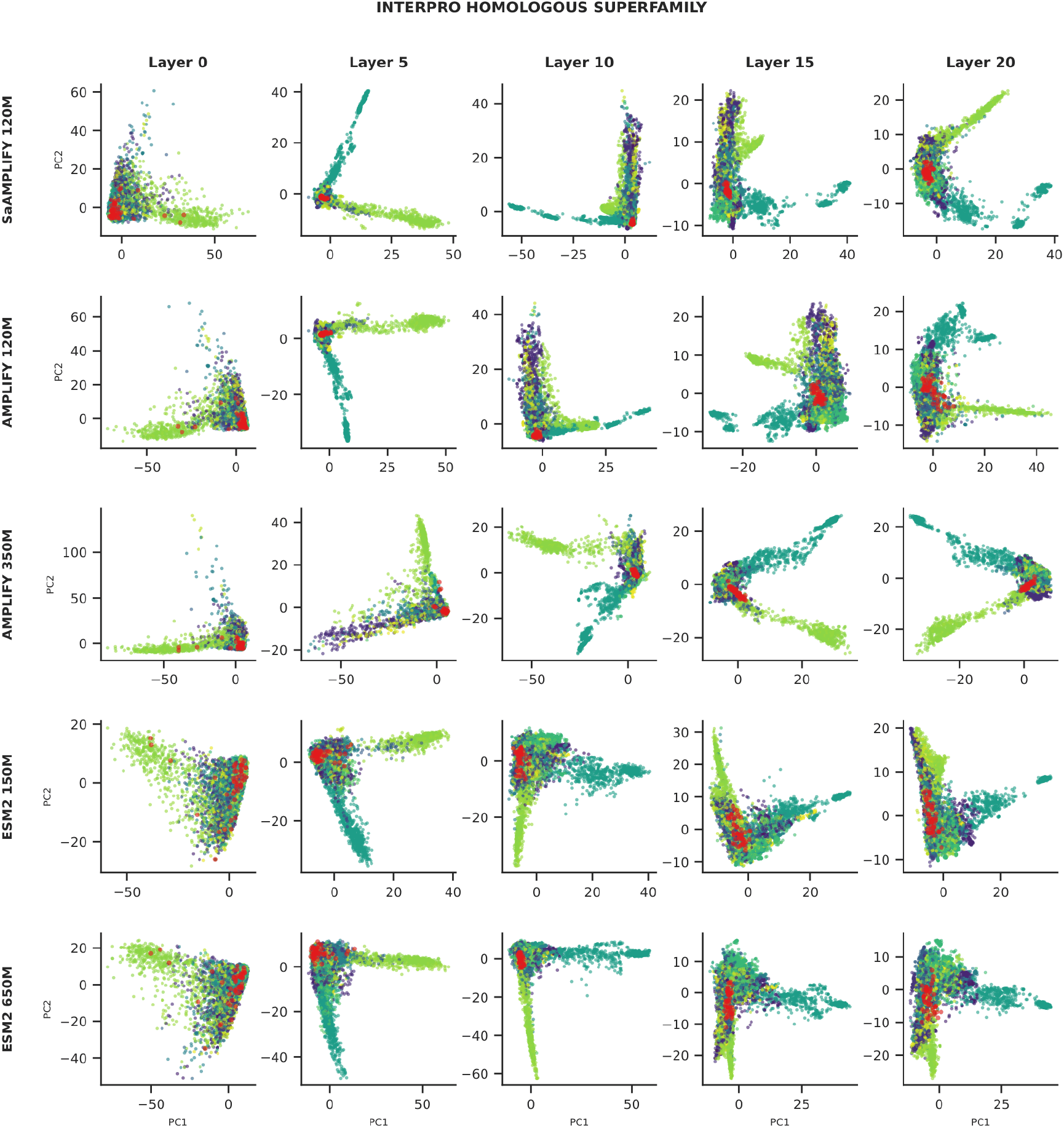
PCA inspection of layerwise embeddings labeled with InterPro Homologous Superfamily annotations shows similarities and differences in organization of groupings. Axis ticks denote 20-unit increments.

**Figure 23.**
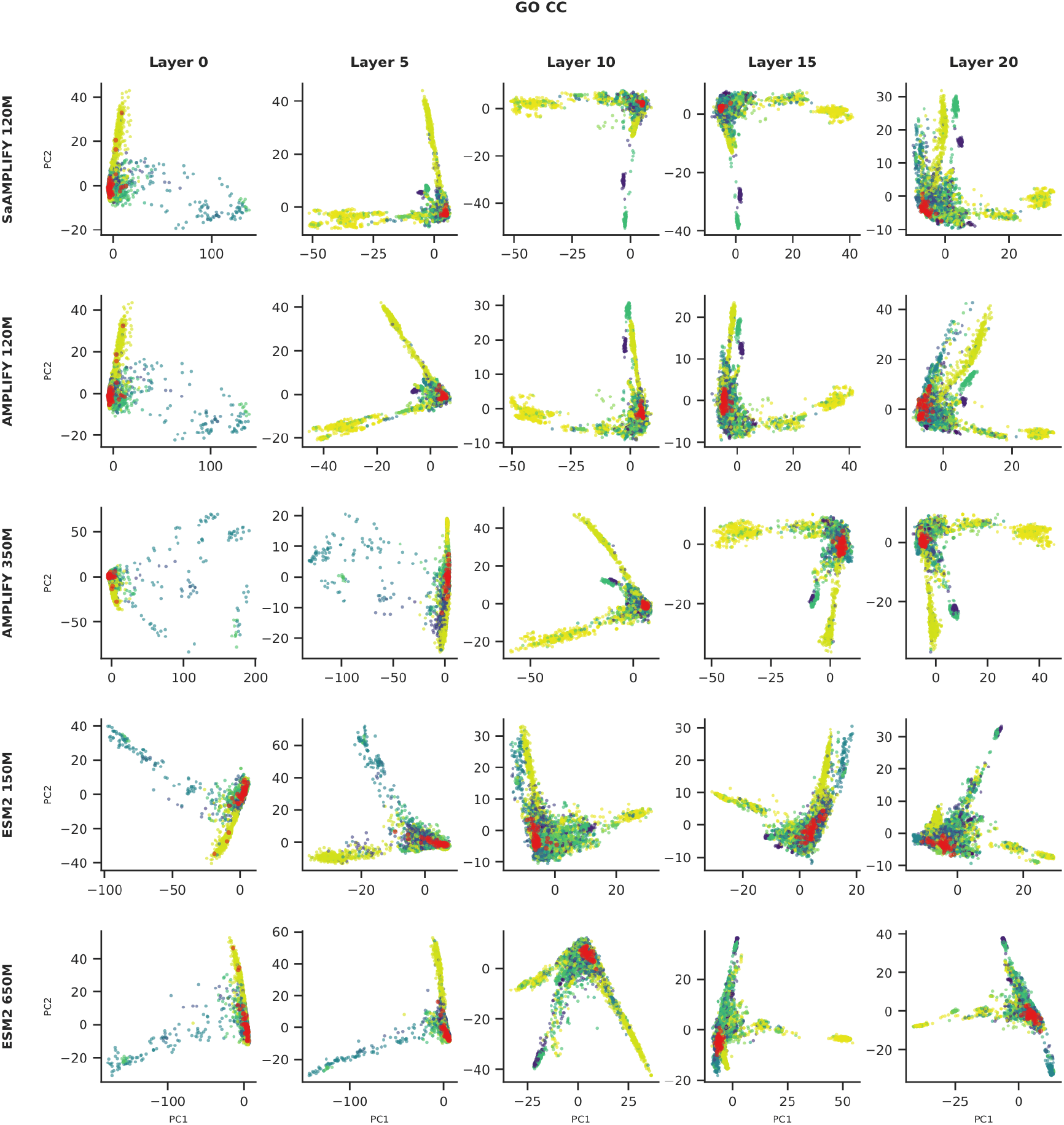
PCA inspection of layerwise embeddings labeled with GO CC annotations shows similarities and differences in organization of groupings. Axis ticks denote 20-unit increments.

**Figure 24.**
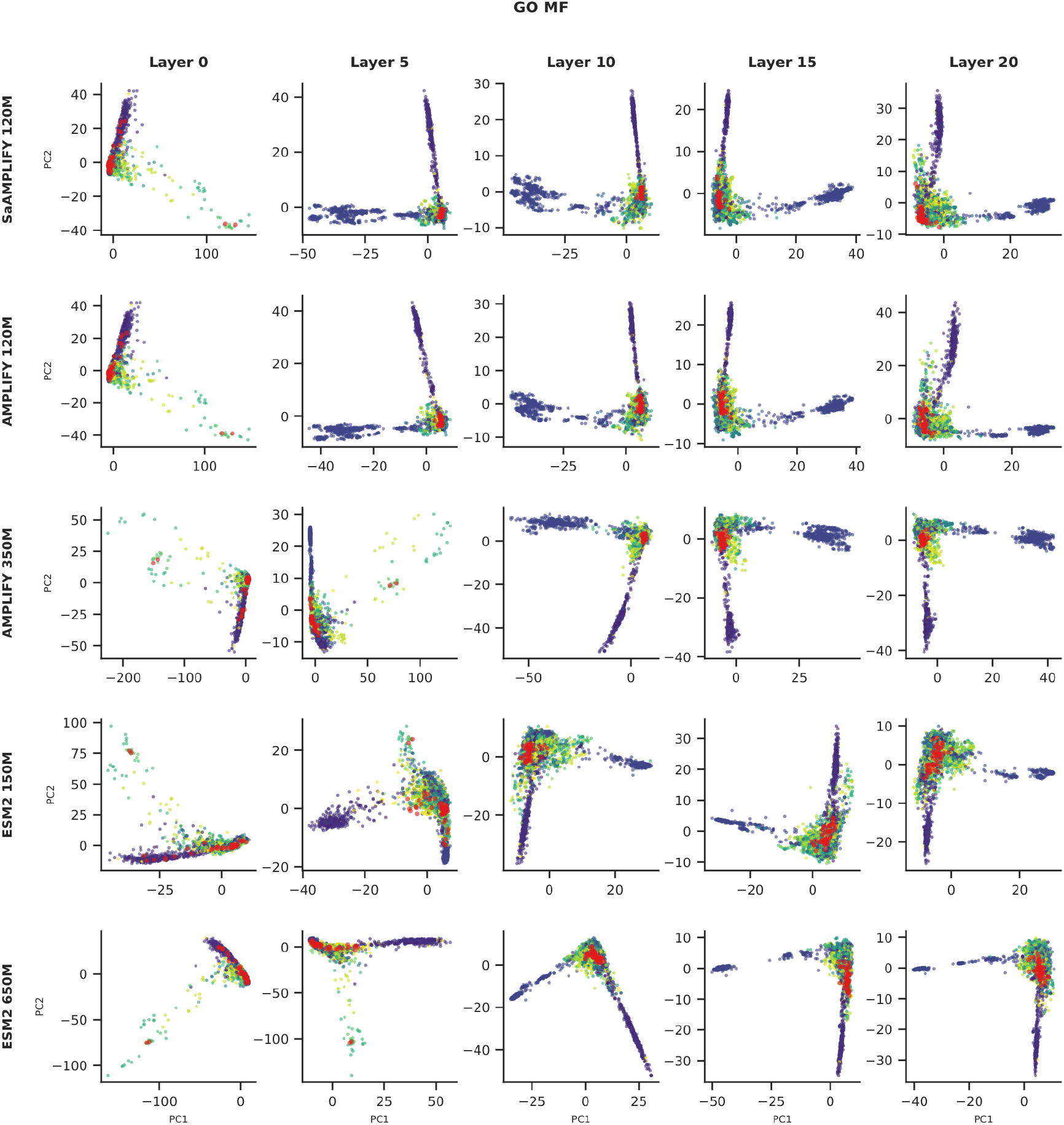
PCA inspection of layerwise embeddings labeled with GO MF annotations shows similarities and differences in organization of groupings. Axis ticks denote 20-unit increments.

**Figure 25.**
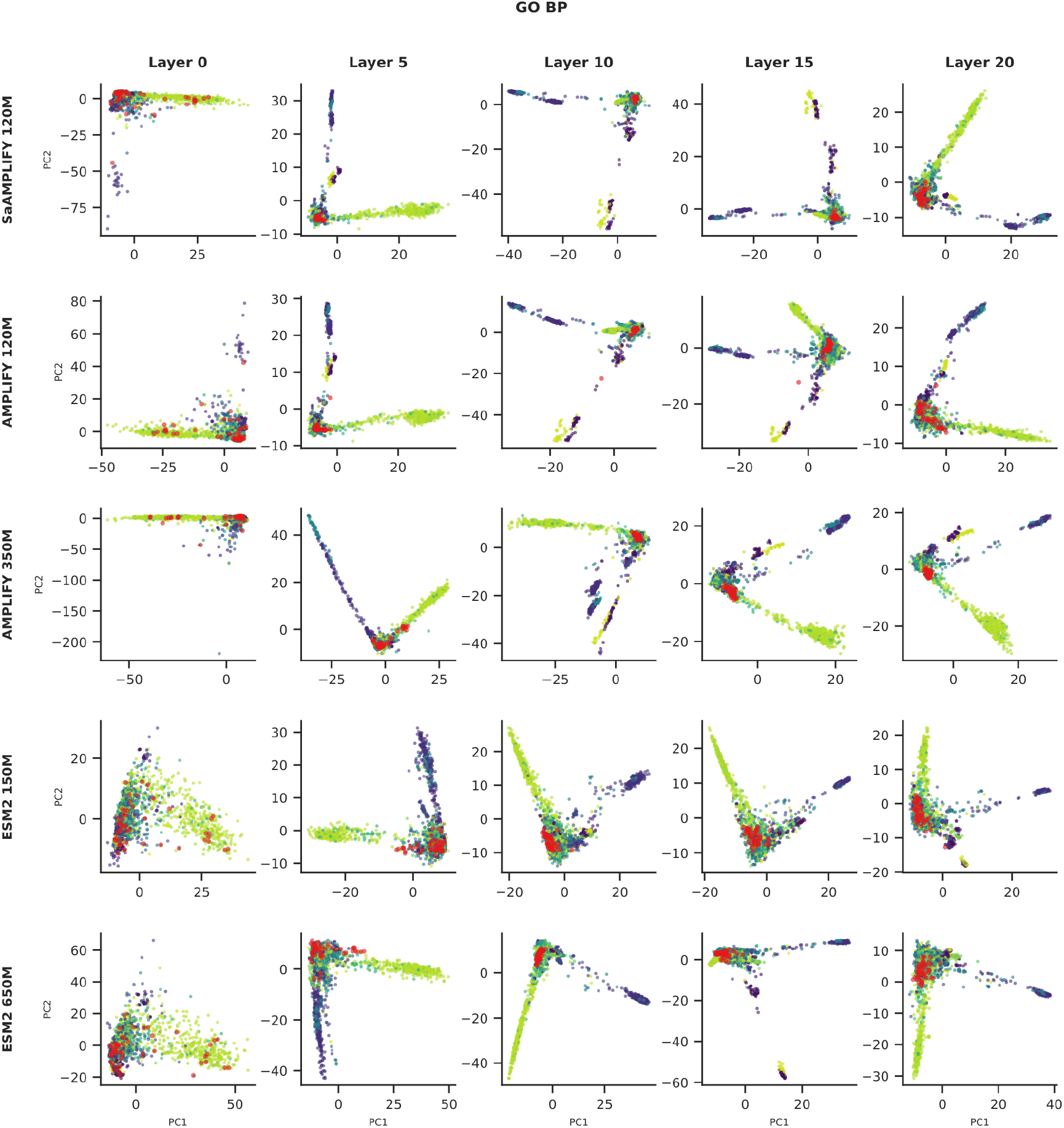
PCA inspection of layerwise embeddings labeled with GO BP annotations shows similarities and differences in organization of groupings. Axis ticks denote 20-unit increments.

**Figure 26.**
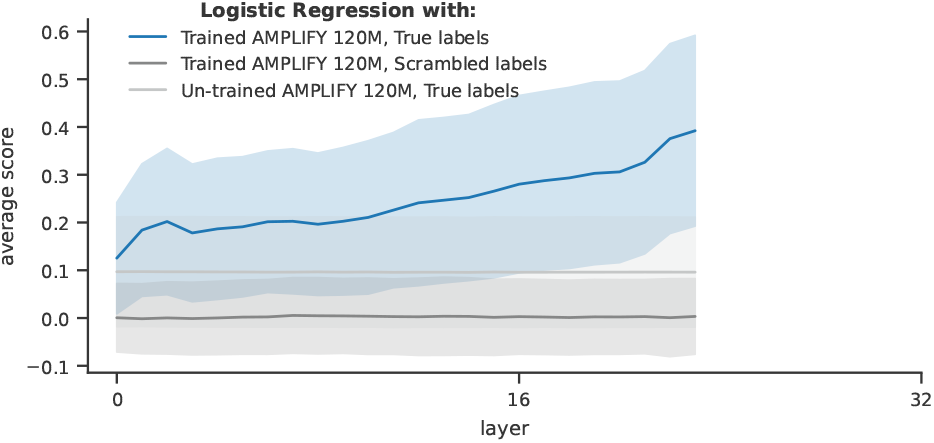
Molecular-biology-inspired manipulations of protein sequences suggest that we can use embeddings to distinguish between theoretically active and inactive forms of a protein. For each randomly-sampled protein (n=1000 samplings), each serine (Ser), threonine (Thr), or tyrosine (Tyr) residue was mutated to alanine (Ala), glycine (Gly), or glutamate (Glu) and a logistic regression classifier trained on a stratified 70:30 split of AMPLIFY 120M embeddings to predict whether a mutant is anticipated to be active (wild-type or Glu at UniProt-annotated phosphosites), inactive (Gly or Ala at annotated phosphosites), or unknown effect (mutations at Ser/Thr/Tyr other than those annotated in UniProt). Average score is the average Matthews Correlation Coefficient among all proteins. Logistic regressors were trained to either predict true labels or scrambled labels from trained AMPLIFY 120M or an un-trained version whose weights and biases were reset to PyTorch defaults. Shaded areas are standard deviation.

**Figure 27.**
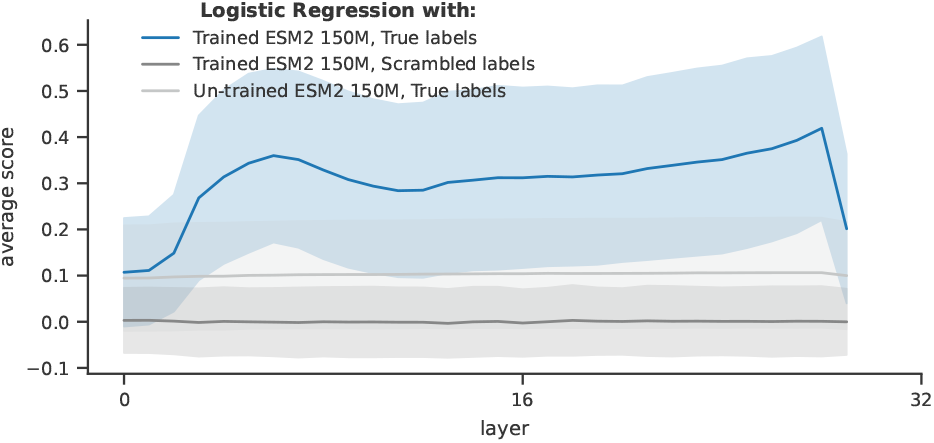
Molecular-biology-inspired manipulations of protein sequences suggest that we can use embeddings to distinguish between theoretically active and inactive forms of a protein. For each randomly-sampled protein (n=1000 samplings), each serine (Ser), threonine (Thr), or tyrosine (Tyr) residue was mutated to alanine (Ala), glycine (Gly), or glutamate (Glu) and a logistic regression classifier trained on a stratified 70:30 split of ESM2 150M embeddings to predict whether a mutant is anticipated to be active (wild-type or Glu at UniProt-annotated phosphosites), inactive (Gly or Ala at annotated phosphosites), or unknown effect (mutations at Ser/Thr/Tyr other than those annotated in UniProt). Average score is the average Matthews Correlation Coefficient among all proteins. Logistic regressors were trained to either predict true labels or scrambled labels from trained ESM2 150M or an un-trained version whose weights and biases were reset to PyTorch defaults. Shaded areas are standard deviation.

**Figure 28.**
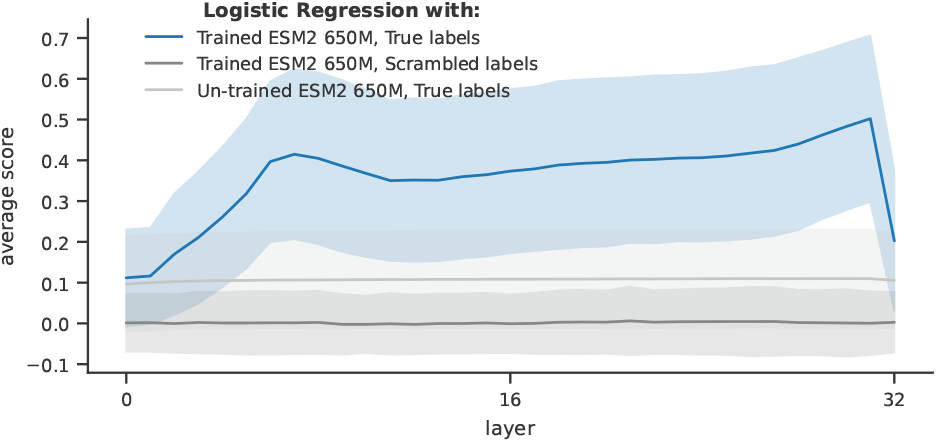
Molecular-biology-inspired manipulations of protein sequences suggest that we can use embeddings to distinguish between theoretically active and inactive forms of a protein. For each randomly-sampled protein (n=1000 samplings), each serine (Ser), threonine (Thr), or tyrosine (Tyr) residue was mutated to alanine (Ala), glycine (Gly), or glutamate (Glu) and a logistic regression classifier trained on a stratified 70:30 split of ESM2 650M embeddings to predict whether a mutant is anticipated to be active (wild-type or Glu at UniProt-annotated phosphosites), inactive (Gly or Ala at annotated phosphosites), or unknown effect (mutations at Ser/Thr/Tyr other than those annotated in UniProt). Average score is the average Matthews Correlation Coefficient among all proteins. Logistic regressors were trained to either predict true labels or scrambled labels from trained ESM2 650M or an un-trained version whose weights and biases were reset to PyTorch defaults. Shaded areas are standard deviation.

## Discussion

Interpretability studies are essential for revealing biases, understanding how design choices impact the information encoded by models, ensuring they do not rely on spurious statistical artifacts, and ultimately encouraging trust and adoption among end users. Accordingly, we systematically investigated how various pLM configurations and families acquire a broad range of overlapping biology concepts, revealing their strengths and limitations.

Our linear probing strategy showed that pLM representations encode a broad range of concepts, capturing different levels of biological complexity. Some of these concepts are more important for a model to learn than others: although predicting molecular weight and other ProtParams was useful for validating our controls, it is not a task for which one would use a pLM in practice. Similarly, programs to predict secondary structure, repeats, and even protein localization are fairly well developed. Otherwise, “which concepts are more important to learn” depends on the scientist and the goal: knowing that pLM representations can predict active or binding sites may be exciting for an industry scientist, while an evolutionary biologist may be more interested in pLMs’ use for remote homology detection through their ability to identify protein domains and families. The hint that pLMs develop sophisticated, nonlinear representations past layer 12 is tantalizing for those who want to understand protein interaction biology, and will hopefully drive further investigation.

Robust controls are essential to all scientific research, and particularly in interpretability studies. In this work, we implemented controls inspired by molecular biology, single-cell perturbation prediction, and first principles. Scrambled sequence controls suggest that the metal-ion-binding and protein-localization datasets obtained from XTrimo-PGLM (BioMap) are of poor quality. This reflects a well-known issue in computational biology: most benchmarks are outdated or poorly designed, and only a subset are truly useful ^9,58,64^. In an attempt to circumvent these limitations, we sourced annotations from UniProt, a highly regarded protein database, and focused on the most-studied proteome: the human proteome. Nevertheless, many proteins remain poorly characterized. Furthermore, biological concepts such as secondary structure, domains, and functions are deeply intertwined. Within our computational constraints, we used subsampling and orthologous approaches where possible to estimate error and increase confidence in our conclusions. While a higher-budget study or one with a narrower focus might yield cleaner results, our study, to our knowledge, examines the broadest range of concepts and models in pLM interpretability while maintaining high layer-wise resolution. Our controls also revealed that, for residue-wise probing, normalization may help signal detection (see Appendix). We believe this finding suggests that residue-wise probes detect relative differences in embedding positions, whereas protein-level probes gain more information from embedding magnitudes.

A key contribution of this study is showing that biological concepts emerge at increasing levels of complexity along pLM layers, and that this emergence is shaped more by pretraining data than by parameter scaling. The superior performance of AMPLIFY models at early layers (Figure 4) and the observed benefits of training on UniRef100 compared to UniRef50 (Figure 6) support this: for several concepts, data diversity shapes how well they are acquired, with additional compute then driving the gains. The inclusion of diverse sequences, such as SCOP domains and antibodies, in AMPLIFY likely provides a richer signal for learning localized structural and functional features earlier in the model architecture. Future research should investigate the impact of pretraining dataset composition and diversity on the concepts examined in this study. A natural extension of this work would be to evaluate other prominent pLM model families using the same approaches to determine whether models with different objectives, such as next-token prediction rather than masked language modeling, change the trends we observed. Furthermore, we observed that pLMs may make complex nonlinear representational adjustments beyond layer 12 that are not fully captured by the current experimental approach. Future research should also investigate the nonlinear encoding of concepts.

A study concurrent to our work investigated whether deep learning models for co-folding learn the physics of protein-ligand interactions ^39^. Similar to our molecular-biology-inspired interventions, the study used adversarial examples based on established physical, chemical, and biological principles, demonstrating the importance of domain expertise in interpretability research. The phosphorylation intervention illustrates the value of pairing pLM interpretability with hypotheses drawn from molecular biology. Future work could expand upon our methodology by implementing similar interventions at the sequence level, such as evaluating whether pLMs can distinguish between evolutionarily plausible domain swapping or correctly identify engineered cleavage sites. We strongly believe that such mechanistic interventions, designed jointly with domain experts, will be the most informative next step for understanding what pLMs encode.

## Methods

### Reproducibility

Random seed was set to 184502 for all data preprocessing and python notebooks. For probe training and evaluation, the random seed was set to 24895.

### Data access and preprocessing

The dataset of proteins from the Homo sapiens proteome with the status “Reviewed” was obtained from UniProt ^18^ at https://www.uniprot.org/uniprotkb on 2025-09-09 by selecting the relevant columns and downloading the results as a TSV file. Since ESM2 was trained on proteins with 1024 or fewer residues and performance declines for longer sequences, all proteins exceeding 1024 residues were excluded, reducing the dataset size from 20,420 to 18,126 proteins. For the analysis of AMPLIFY checkpoints, as the model had not yet undergone context extension, the dataset was further restricted to proteins with 512 or fewer residues. BioMap datasets ^12^ (see Table 1) were obtained through HuggingFace.

### Protein analysis with BioPython

The ProteinAnalysis tool from BioPython ^16^ was used to calculate the following properties for each sequence: the percentage of each amino acid; the fraction of amino acids found in Helix (V, I, Y, F, W, and L), Turn (N, P, G, and S), and Sheet (E, M, A, and L); isoelectric point; charge at pH 4.7, 7.2, and 8.0, which correspond to lysosomal, cytosolic, and mitochondrial pH ^10^; GRAVY, the grand average of hydropathy ^34^; and instability index ^27^. Noncanonical amino acids were replaced with loosely analogous ones: U with C, O with L, B with N, Z with Q, J with L, and X with G. This substitution affected 36 positions across 25 proteins. The protein-level probing task for this dataset was to predict the calculated value of each of these properties per protein as a regression task.

### Parsing and formatting UniProt information

Annotations in Gene Ontology (biological process), Gene Ontology (cellular component), Gene Ontology (molecular function), and InterPro columns were converted to one-hot matrices. Mappings from InterPro identifiers to entry type and name were downloaded from InterPro on 2025-09-16. Separate matrices for each InterPro entry type (Homologous Superfamily, Family, Domain, Active Site, Binding Site, Post-translational Modification Site, Conserved site, Repeat) (see InterPro entry types here) were constructed. There were too few unique domains for the InterPro PTM category, so we dropped it from our analyses. The protein-level probing task for these datasets was to predict the presence or absence of each distinct annotation in a protein (prediction of a one-hot encoding).

### Residue-level annotations

Uniprot annotations were parsed from the TSV file into residue-wise annotations. Conceptually similar categories were grouped together into individual datasets, with Secondary Structure being composed of Helix, Strand, and Turn annotations; Post-Translational Modification of Carbohydrate, Disulfide, Modified Residue, and Lipid annotations; and Peptide of Propeptide, Transit Peptide, or Signal Peptide annotations. For Topology, we parsed the note field, giving annotations for whether a residue is expected to be exposed to cytoplasm or environments like nuclear, vacuolar, or mitochondrial matrix compartments. The amino-acid-level probing task for these datasets was to predict the category to which an individual residue belonged, e.g., Helix, Strand, Turn, or None (see Table 2 for the full set of classes per residue-level dataset).

### Train-validation-test splits

We used MMseqs2^54,55^ easy-cluster to group protein sequences at increasing levels of minimum sequence identity (min-seq-id), only clustering sequences that have a sequence length overlap greater than 80% of the target sequence (-c 0.8, --cov-mode 1). Test sets were assigned by randomly selecting proteins with different representative members at a min-seq-id of 0.20; at this clustering level, a protein in the test set should not be more than 20% identical to one in the training set. Validation sets were constructed by subsampling the remaining proteins based on representative membership, with a minimum sequence identity of 0.5 to balance generalization and training evaluation. Proteins with InterPro Active Site, Binding Site, Conserved Site, and Repeat annotations were split at a 0.30 min-seq-id, since the individual InterPro domains are fine-grained and family-specific enough that a more strict split would largely prevent a domain from being present in both train and test sets.

### Training and evaluating linear probes

For ESM2 models, the embedding layer at the model’s input was treated as external and was neither included in the depth calculation nor used as a layer to probe. Linear probes consisted of a single PyTorch Linear layer with input dimensions equal to those of the pLM’s hidden dimension, and output dimensions equal to the number of targets. Probes were trained with early stopping for 50 epochs with an AdamW optimiser with constant learning_rate 1e-3, weight_decay 1e-4, adam_epsilon 1e-8, and adam_betas [0.9, 0.999]. After training, the best performing checkpoint is loaded, and the test step scored the embeddings from a pLM or its “un-trained” control and a linear probe or its “un-trained” control. Controls are given in Table 3:

Target labels per protein were changed depending on the control and the task type (regression, binary, or multiclass classification) and task level (protein or residue-level) as shown in Table 4.

7 different random subsets (taking 50% of predictions each time) of the test set were scored to get an estimate of test set variability. Because of the generally high sparsity of amino-acid-level annotations, we only calculated elementwise metrics for protein-level tasks. Although we calculated a range of score metrics, we primarily report Matthews Correlation Coefficient:

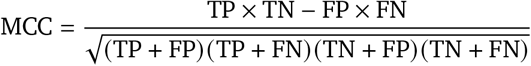

TP (true positive) is the number of true positive predictions (positive cases that were correctly identified as positive). TN (true negative) is the number of true negative predictions (negative cases that were correctly identified as negative). FP (false positive) is the number of false positive predictions (negative cases that were incorrectly identified as positive). FN (false negative) is the number of false negative predictions (positive cases that were incorrectly identified as negative). The MCC accounts for class imbalance and generates a high score in its interval [-1,1] only if the classifier scored a high value for all the four basic rates of the confusion matrix: sensitivity, specificity, precision, and negative predictive value and has been suggested to be more suitable than F1 score or area under the ROC curve (AUC) for binary classification problems ^14,15^. MCC can also be applied to a multiclass situation:

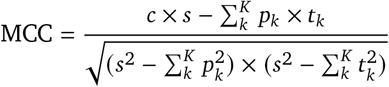

Where

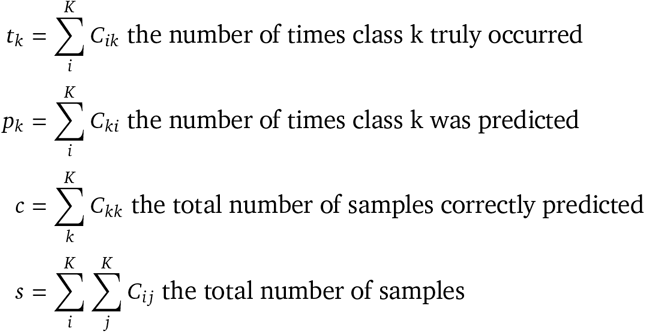

### Molecular biology intervention experiments

#### Phosphorylation

We parsed notes from the Modified Residue field of our UniProt TSV, which indicates the kind of residue modified. After filtering for annotations for phosphoserine, phosphothreonine, or phosphotyrosine, we performed scanning mutagenesis. For each protein, we changed each serine, threonine, or tyrosine to an alanine, a glycine (both predicted to be either neutral or deleterious to protein function unless the mutation is made at a phosphosite, in which case it is predicted to be inactivating), or a glutamate (predicted to be hyper-activating if the residue is a phosphosite), and labeled it with the predicted outcome Active/Inactive/Unknown. We annotated wild-type sequences as Active. We tokenized mutated and wild-type sequences, fed them into pLMs, and extracted raw embeddings for each layer. We constructed the un-trained pLM control the same way as for the linear probing controls. We split embeddings and labels 70:30 into train and test sets and trained logistic regression models from scikit-learn with default hyperparameters for 350 iterations to predict whether a mutant is active, inactive, or of unknown effect. As a control, we shuffled labels after splitting to ensure that the logistic regressors do not learn spurious mappings between embeddings and annotations. We repeated this process 1000 times with randomly-selected proteins that have annotated phosphosites in UniProt. We used a pairwise Wilcoxon signed-rank test to evaluate whether there were differences between logistic regressor scores for true labels vs. scrambled labels, per layer, with a Bonferroni post-hoc test to correct for multiple comparisons between layers.

## Acknowledgments

This research was enabled in part by compute resources provided by Mila (mila.quebec). We thank Lola Le Breton for providing the intermediate AMPLIFY checkpoints and preparing them for Hugging Face. Generative AI tools were used to review and proofread the manuscript; all scientific content, analyses, and conclusions are the authors’ own.

## Author contributions

Conceptualization: Shawn T. Whitfield and Quentin Fournier. Data curation: Shawn T. Whitfield. Formal analysis: Shawn T. Whitfield. Funding acquisition: Christopher J. Langmead and Quentin Fournier. Investigation: Shawn T. Whitfield. Methodology: Shawn T. Whitfield, Dhanya Sridhar, and Quentin Fournier. Software: Shawn T. Whitfield and Tom Marty. Supervision: Dhanya Sridhar and Quentin Fournier. Visualization: Shawn T. Whitfield. Writing - original draft: Shawn T. Whitfield. Writing - review & editing: Shawn T. Whitfield, Tom Marty, Robert M. Vernon, Christopher J. Langmead, Dhanya Sridhar, and Quentin Fournier.

## Data availability

The curated datasets, linear probing outputs, intervention results, and source data underlying all figures are available on Figshare (10.6084/m9.figshare.31169506). Reviewed *Homo sapiens* annotations were obtained from UniProt (https://www.uniprot.org/, accessed 2025-09-09). InterPro entry-type mappings were obtained from https://www.ebi.ac.uk/interpro/download/ (accessed 2025-09-16). BioMap datasets are available through HuggingFace (https://huggingface.co/biomap-research/datasets).

## Code availability

All code used in this study is available on GitHub at https://github.com/chandar-lab/AMPLIFY/tree/interpretability under the MIT license.

## Additional information

Correspondence and requests for materials should be addressed to Quentin Fournier (quentin.fournier@ mila.quebec).

## A. Supplementary information

## References

1 Etowah Adams, Liam Bai, Minji Lee, Yiyang Yu, and Mohammed AlQuraishi. From Mechanistic Interpretability to Mechanistic Biology: Training, Evaluating, and Interpreting Sparse Autoencoders on Protein Language Models, June 2025. URL https://www.biorxiv.org/content/10.1101/2025.02.06.636901v2.

2 Antonina Andreeva, Dave Howorth, Cyrus Chothia, Eugene Kulesha, and Alexey G. Murzin. SCOP2 prototype: a new approach to protein structure mining. Nucleic Acids Research, 42(D1):D310–D314, January 2014. ISSN 0305-1048. doi: 10.1093/nar/gkt1242. URL 10.1093/nar/gkt1242.

3 Antonina Andreeva, Eugene Kulesha, Julian Gough, and Alexey G Murzin. The SCOP database in 2020: expanded classification of representative family and superfamily domains of known protein structures. Nucleic Acids Research, 48(D1):D376–D382, January 2020. ISSN 0305-1048. doi: 10.1093/nar/gkz1064. URL 10.1093/nar/gkz1064.

4 Christian B. Anfinsen. Principles that Govern the Folding of Protein Chains. Science, 181(4096):223–230, July 1973. doi: 10.1126/science.181.4096.223. URL https://www.science.org/doi/10.1126/science.181.4096.223.

5 Michael Ashburner, Catherine A. Ball, Judith A. Blake, David Botstein, Heather Butler, J. Michael Cherry, Allan P. Davis, Kara Dolinski, Selina S. Dwight, Janan T. Eppig, Midori A. Harris, David P. Hill, Laurie Issel-Tarver, Andrew Kasarskis, Suzanna Lewis, John C. Matese, Joel E. Richardson, Martin Ringwald, Gerald M. Rubin, and Gavin Sherlock. Gene Ontology: tool for the unification of biology. Nature genetics, 25(1):25–29, May 2000. ISSN 1061-4036. doi: 10.1038/75556. URL https://pmc.ncbi.nlm.nih.gov/articles/PMC3037419/.

6 Arjun Banerjee, David Martinez, Camille Dang, and Ethan Tam. Automated Neuron Labelling Enables Generative Steering and Interpretability in Protein Language Models, July 2025. URL http://arxiv.org/abs/2507.06458.

7 Thomas Bikias, Evangelos Stamkopoulos, and Sai T Reddy. PLMFit: benchmarking transfer learning with protein language models for protein engineering. Briefings in Bioinformatics, 26(4):bbaf381, July 2025. ISSN 1477-4054. doi: 10.1093/bib/bbaf381. URL 10.1093/bib/bbaf381.

8 Matthias Blum, Antonina Andreeva, Laise Cavalcanti Florentino, Sara Rocio Chuguransky, Tiago Grego, Emma Hobbs, Beatriz Lazaro Pinto, Ailsa Orr, Typhaine Paysan-Lafosse, Irina Ponamareva, Gustavo A Salazar, Nicola Bordin, Peer Bork, Alan Bridge, Lucy Colwell, Julian Gough, Daniel H Haft, Ivica Letunic, Felipe Llinares-López, Aron Marchler-Bauer, Laetitia Meng-Papaxanthos, Huaiyu Mi, Darren A Natale, Christine A Orengo, Arun P Pandurangan, Damiano Piovesan, Catherine Rivoire, Christian J A Sigrist, Narmada Thanki, Françoise Thibaud-Nissen, Paul D Thomas, Silvio C E Tosatto, Cathy H Wu, and Alex Bateman. InterPro: the protein sequence classification resource in 2025. Nucleic Acids Research, 53(D1):D444–D456, January 2025. ISSN 1362-4962. doi: 10.1093/nar/gkae1082. URL 10.1093/nar/gkae1082.

9 Yue Cao, Lijia Yu, Marni Torkel, Sanghyun Kim, Yingxin Lin, Pengyi Yang, Terence P Speed, Shila Ghazanfar, and Jean Yee Hwa Yang. The current landscape and emerging challenges of benchmarking single-cell methods. Briefings in Bioinformatics, 26(5):bbaf380, September 2025. ISSN 1477-4054. doi: 10.1093/bib/bbaf380. URL 10.1093/bib/bbaf380.

10 Joseph R. Casey, Sergio Grinstein, and John Orlowski. Sensors and regulators of intracellular pH. Nature Reviews Molecular Cell Biology, 11(1):50–61, January 2010. ISSN 1471-0080. doi: 10.1038/nrm2820. URL https://www.nature.com/articles/nrm2820.

11 Tara Chari and Lior Pachter. The specious art of single-cell genomics. PLOS Computational Biology, 19(8):e1011288, August 2023. ISSN 1553-7358. doi: 10.1371/journal.pcbi.1011288. URL https://journals.plos.org/ploscompbiol/article?id=10.1371/journal.pcbi.1011288.

12 Bo Chen, Xingyi Cheng, Pan Li, Yangli-ao Geng, Jing Gong, Shen Li, Zhilei Bei, Xu Tan, Boyan Wang, Xin Zeng, Chiming Liu, Aohan Zeng, Yuxiao Dong, Jie Tang, and Le Song. xTrimoPGLM: Unified 100B-Scale Pre-trained Transformer for Deciphering the Language of Protein, December 2024. URL http://arxiv.org/abs/2401.06199. arXiv:2401.06199 q-bio].

13 Can Chen, David Heurtel-Depeiges, Robert M. Vernon, Christopher James Langmead, Yoshua Bengio, and Quentin Fournier. Structure-Aligned Protein Language Model, December 2025. URL http://arxiv.org/abs/2505.16896. arXiv:2505.16896 cs].

14 Davide Chicco and Giuseppe Jurman. The advantages of the Matthews correlation coefficient (MCC) over F1 score and accuracy in binary classification evaluation. BMC genomics, 21(1):6, January 2020. ISSN 1471-2164. doi: 10.1186/s12864-019-6413-7.

15 Davide Chicco and Giuseppe Jurman. The Matthews correlation coefficient (MCC) should replace the ROC AUC as the standard metric for assessing binary classification. BioData Mining, 16(1):4, February 2023. ISSN 1756-0381. doi: 10.1186/s13040-023-00322-4.

16 Peter J. A. Cock, Tiago Antao, Jeffrey T. Chang, Brad A. Chapman, Cymon J. Cox, Andrew Dalke, Iddo Friedberg, Thomas Hamelryck, Frank Kauff, Bartek Wilczynski, and Michiel J. L. de Hoon. Biopython: freely available Python tools for computational molecular biology and bioinformatics. Bioinformatics, 25(11):1422–1423, June 2009. ISSN 1367-4803. doi: 10.1093/bioinformatics/btp163. URL 10.1093/bioinformatics/btp163.

17 The Gene Ontology Consortium. The Gene Ontology knowledgebase in 2026. Nucleic Acids Research, 54(D1):D1779–D1792, January 2026. ISSN 1362-4962. doi: 10.1093/nar/gkaf1292. URL 10.1093/nar/gkaf1292.

18 The UniProt Consortium. UniProt: the Universal Protein Knowledgebase in 2025. Nucleic Acids Research, 53(D1):D609–D617, January 2025. ISSN 1362-4962. doi: 10.1093/nar/gkae1010. URL 10.1093/nar/gkae1010.

19 Tri Dao, Daniel Y. Fu, Stefano Ermon, Atri Rudra, and Christopher Ré. FlashAttention: Fast and Memory-Efficient Exact Attention with IO-Awareness, June 2022. URL http://arxiv.org/abs/2205.14135. arXiv:2205.14135 cs].

20 Nicki Skafte Detlefsen, Søren Hauberg, and Wouter Boomsma. Learning meaningful representations of protein sequences. Nature Communications, 13(1):1914, April 2022. ISSN 2041-1723. doi: 10.1038/s41467-022-29443-w. URL https://www.nature.com/articles/s41467-022-29443-w.

21 Nicolas Deutschmann, Aurelien Pelissier, Anna Weber, Shuaijun Gao, Jasmina Bogojeska, and María Rodríguez Martínez. Do domain-specific protein language models outperform general models on immunology-related tasks? ImmunoInformatics, 14, June 2024. ISSN 2667-1190. doi: 10.1016/j.immuno.2024.100036. URL https://www.immunoinformaticsjournal.com/article/S2667-1190%2824%2900006-5/fulltext#tbl2.

22 Jacob Devlin, Ming-Wei Chang, Kenton Lee, and Kristina Toutanova. BERT: Pre-training of Deep Bidirectional Transformers for Language Understanding, May 2019. URL http://arxiv.org/abs/1810.04805. arXiv:1810.04805 cs].

23 Ahmed Elnaggar, Michael Heinzinger, Christian Dallago, Ghalia Rehawi, Yu Wang, Llion Jones, Tom Gibbs, Tamas Feher, Christoph Angerer, Martin Steinegger, Debsindhu Bhowmik, and Burkhard Rost. ProtTrans: Towards Cracking the Language of Life’s Code Through Self-Supervised Learning, July 2020. URL http://biorxiv.org/lookup/doi/10.1101/2020.07.12.199554.

24 Quentin Fournier, Robert M. Vernon, Almer van der Sloot, Benjamin Schulz, Sarath Chandar, and Christopher James Langmead. Protein Language Models: Is Scaling Necessary?, January 2026. URL https://www.biorxiv.org/content/10.1101/2024.09.23.614603v2. ISSN: 2692-8205 Pages: 2024.09.23.614603 Section: New Results.

25 Edith Natalia Villegas Garcia and Alessio Ansuini. Interpreting and Steering Protein Language Models through Sparse Autoencoders, February 2025. URL http://arxiv.org/abs/2502.09135.

26 Onkar Gujral, Mihir Bafna, Eric Alm, and Bonnie Berger. Sparse autoencoders uncover biologically interpretable features in protein language model representations. Proceedings of the National Academy of Sciences, 122(34):e2506316122, August 2025. doi: 10.1073/pnas.2506316122. URL https://www.pnas.org/doi/abs/10.1073/pnas.2506316122. Company: National Academy of Sciences Distributor: National Academy of Sciences Institution: National Academy of Sciences Label: National Academy of Sciences.

27 K. Guruprasad, B. V. Reddy, and M. W. Pandit. Correlation between stability of a protein and its dipeptide composition: a novel approach for predicting in vivo stability of a protein from its primary sequence. Protein Engineering, 4(2):155–161, December 1990. ISSN 0269-2139. doi: 10.1093/protein/4.2.155.

28 Michael Heinzinger, Ahmed Elnaggar, Yu Wang, Christian Dallago, Dmitrii Nechaev, Florian Matthes, and Burkhard Rost. Modeling aspects of the language of life through transfer-learning protein sequences. BMC Bioinformatics, 20(1):723, December 2019. ISSN 1471-2105. doi: 10.1186/s12859-019-3220-8. URL 10.1186/s12859-019-3220-8.

29 Haiyang Huang, Yingfan Wang, Cynthia Rudin, and Edward P. Browne. Towards a comprehensive evaluation of dimension reduction methods for transcriptomic data visualization. Communications Biology, 5:719, July 2022. ISSN 2399-3642. doi: 10.1038/s42003-022-03628-x. URL https://pmc.ncbi.nlm.nih.gov/articles/PMC9296444/.

30 Eric Kernfeld, Yunxiao Yang, Joshua S. Weinstock, Alexis Battle, and Patrick Cahan. A comparison of computational methods for expression forecasting. Genome Biology, 26(1):388, November 2025. ISSN 1474-760X. doi: 10.1186/s13059-025-03840-y. URL 10.1186/s13059-025-03840-y.

31 Henry R. Kilgore, Itamar Chinn, Peter G. Mikhael, Ilan Mitnikov, Catherine Van Dongen, Guy Zylberberg, Lena Afeyan, Salman F. Banani, Susana Wilson-Hawken, Tong Ihn Lee, Regina Barzilay, and Richard A. Young. Protein codes promote selective subcellular compartmentalization. Science, 387(6738):1095–1101, March 2025. doi: 10.1126/science.adq2634. URL https://www.science.org/doi/10.1126/science.adq2634.

32 Aleksandr Kovaltsuk, Jinwoo Leem, Sebastian Kelm, James Snowden, Charlotte M Deane, and Konrad Krawczyk. Observed Antibody Space: A Resource for Data Mining Next-Generation Sequencing of Antibody Repertoires. The Journal of Immunology, 201(8):2502–2509, October 2018. ISSN 0022-1767. doi: 10.4049/jimmunol.1800708. URL 10.4049/jimmunol.1800708.

33 Kyohei Koyama, Kosuke Hashimoto, Chioko Nagao, and Kenji Mizuguchi. Attention network for predicting T-cell receptor–peptide binding can associate attention with interpretable protein structural properties. Frontiers in Bioinformatics, 3, December 2023. ISSN 2673-7647. doi: 10.3389/fbinf.2023.1274599. URL https://www.frontiersin.org/journals/bioinformatics/articles/10.3389/fbinf.2023.1274599/full.

34 J. Kyte and R. F. Doolittle. A simple method for displaying the hydropathic character of a protein. Journal of Molecular Biology, 157(1):105–132, May 1982. ISSN 0022-2836. doi: 10.1016/0022-2836(82)90515-0.

35 Francesca-Zhoufan Li, Ava P. Amini, Yisong Yue, Kevin K. Yang, and Alex X. Lu. Feature Reuse and Scaling: Understanding Transfer Learning with Protein Language Models, February 2024. URL https://www.biorxiv.org/content/10.1101/2024.02.05.578959v2. Pages: 2024.02.05.578959 Section: New Results.

36 Zeming Lin, Halil Akin, Roshan Rao, Brian Hie, Zhongkai Zhu, Wenting Lu, Nikita Smetanin, Robert Verkuil, Ori Kabeli, Yaniv Shmueli, Allan dos Santos Costa, Maryam Fazel-Zarandi, Tom Sercu, Salvatore Candido, and Alexander Rives. Evolutionary-scale prediction of atomic-level protein structure with a language model. Science, 379(6637):1123–1130, March 2023. doi: 10.1126/science.ade2574. URL https://www.science.org/doi/10.1126/science.ade2574.

37 Maria Littmann, Michael Heinzinger, Christian Dallago, Tobias Olenyi, and Burkhard Rost. Embeddings from deep learning transfer GO annotations beyond homology. Scientific Reports, 11(1):1160, January 2021. ISSN 2045-2322. doi: 10.1038/s41598-020-80786-0. URL https://www.nature.com/articles/s41598-020-80786-0.

38 Céline Marquet, Michael Heinzinger, Tobias Olenyi, Christian Dallago, Kyra Erckert, Michael Bernhofer, Dmitrii Nechaev, and Burkhard Rost. Embeddings from protein language models predict conservation and variant effects. Human Genetics, 141(10):1629–1647, October 2022. ISSN 1432-1203. doi: 10.1007/s00439-021-02411-y. URL 10.1007/s00439-021-02411-y.

39 Matthew R. Masters, Amr H. Mahmoud, and Markus A. Lill. Investigating whether deep learning models for co-folding learn the physics of protein-ligand interactions. Nature Communications, 16(1):8854, October 2025. ISSN 2041-1723. doi: 10.1038/s41467-025-63947-5. URL https://www.nature.com/articles/s41467-025-63947-5.

40 Mengren, Liu, Yixiang Zhang, Yiming, and Zhang. Exploring Protein Language Model Architecture-Induced Biases for Antibody Comprehension, December 2025. URL http://arxiv.org/abs/2512.09894. arXiv:2512.09894 cs version: 1.

41 Jatin Nainani, Bryn Marie Reimer, Connor Watts, David Jensen, and Anna G. Green. Mechanistic evidence that motif-gated domain recognition drives contact prediction in protein language models, August 2025. URL https://www.biorxiv.org/content/10.1101/2025.08.22.671739v1. ISSN: 2692-8205 Pages: 2025.08.22.671739 Section: New Results.

42 Gowri Nayar, Alp Tartici, and Russ B. Altman. Paying Attention to Attention: High Attention Sites as Indicators of Protein Family and Function in Language Models, December 2024. URL https://www.biorxiv.org/content/10.1101/2024.12.13.628435v1. Pages: 2024.12.13.628435 Section: New Results.

43 Divya Nori, Shivali Singireddy, and Marina Ten Have. Identification of Knowledge Neurons in Protein Language Models, December 2023. URL http://arxiv.org/abs/2312.10770. arXiv:2312.10770 cs].

44 Krzysztof Odrzywolek, Zuzanna Karwowska, Jan Majta, Aleksander Byrski, Kaja Milanowska-Zabel, and Tomasz Kosciolek. Deep embeddings to comprehend and visualize microbiome protein space. Scientific Reports, 12(1):10332, June 2022. ISSN 2045-2322. doi: 10.1038/s41598-022-14055-7. URL https://www.nature.com/articles/s41598-022-14055-7.

45 Marius Thrane Ødum, Felix Teufel, Vineet Thumuluri, José Juan Almagro Armenteros, Alexander Rosenberg Johansen, Ole Winther, and Henrik Nielsen. DeepLoc 2.1: multi-label membrane protein type prediction using protein language models. Nucleic Acids Research, 52(W1):W215–W220, July 2024. ISSN 0305-1048. doi: 10.1093/nar/gkae237. URL 10.1093/nar/gkae237.

46 Tobias H. Olsen, Fergus Boyles, and Charlotte M. Deane. Observed Antibody Space: A diverse database of cleaned, annotated, and translated unpaired and paired antibody sequences. Protein Science, 31(1): 141–146, 2022. ISSN 1469-896X. doi: 10.1002/pro.4205. URL https://onlinelibrary.wiley.com/doi/abs/10.1002/pro.4205. _eprint: https://onlinelibrary.wiley.com/doi/pdf/10.1002/pro.4205.

47 Kiho Park, Yo Joong Choe, and Victor Veitch. The linear representation hypothesis and the geometry of large language models. In Proceedings of the 41st International Conference on Machine Learning, ICML’24. JMLR.org, 2024.

48 Nithin Parsan, David J. Yang, and John J. Yang. Towards Interpretable Protein Structure Prediction with Sparse Autoencoders, March 2025. URL http://arxiv.org/abs/2503.08764.

49 Pia Francesca Rissom, Paulo Yanez Sarmiento, Jordan Safer, Connor W. Coley, Bernhard Y. Renard, Henrike O. Heyne, and Sumaiya Iqbal. Decoding protein language models: insights from embedding space analysis, February 2025. URL https://www.biorxiv.org/content/10.1101/2024.06.21.600139v2. Pages: 2024.06.21.600139 Section: New Results.

50 Alexander Rives, Joshua Meier, Tom Sercu, Siddharth Goyal, Zeming Lin, Jason Liu, Demi Guo, Myle Ott, C. Lawrence Zitnick, Jerry Ma, and Rob Fergus. Biological structure and function emerge from scaling unsupervised learning to 250 million protein sequences. Proceedings of the National Academy of Sciences, 118(15):e2016239118, April 2021. doi: 10.1073/pnas.2016239118. URL https://www.pnas.org/doi/full/10.1073/pnas.2016239118.

51 Tobias Senoner, Tobias Olenyi, Michael Heinzinger, Anton Spannagl, George Bouras, Burkhard Rost, and Ivan Koludarov. ProtSpace: A Tool for Visualizing Protein Space. Journal of Molecular Biology, 437(15):168940, August 2025. ISSN 0022-2836. doi: 10.1016/j.jmb.2025.168940. URL https://www.sciencedirect.com/science/article/pii/S0022283625000063.

52 Jake Silberg, Elana Simon, and James Zou. Towards functional annotation with latent protein language model features, October 2025. URL https://www.biorxiv.org/content/10.1101/2025.10.02. 680154v1. ISSN: 2692-8205 Pages: 2025.10.02.680154 Section: New Results.

53 Elana Simon and James Zou. InterPLM: discovering interpretable features in protein language models via sparse autoencoders. Nature Methods, 22(10):2107–2117, October 2025. ISSN 1548-7105. doi: 10.1038/s41592-025-02836-7. URL https://www.nature.com/articles/s41592-025-02836-7.

54 Martin Steinegger and Johannes Söding. MMseqs2 enables sensitive protein sequence searching for the analysis of massive data sets. Nature Biotechnology, 35(11):1026–1028, November 2017. ISSN 1546-1696. doi: 10.1038/nbt.3988. URL https://www.nature.com/articles/nbt.3988.

55 Martin Steinegger and Johannes Söding. Clustering huge protein sequence sets in linear time. Nature Communications, 9(1):2542, June 2018. ISSN 2041-1723. doi: 10.1038/s41467-018-04964-5. URL https://www.nature.com/articles/s41467-018-04964-5.

56 Hannes Stärk, Christian Dallago, Michael Heinzinger, and Burkhard Rost. Light attention predicts protein location from the language of life. Bioinformatics Advances, 1(1):vbab035, 01 2021. ISSN 2635-0041. doi: 10.1093/bioadv/vbab035. URL 10.1093/bioadv/vbab035.

57 Jianlin Su, Yu Lu, Shengfeng Pan, Ahmed Murtadha, Bo Wen, and Yunfeng Liu. RoFormer: Enhanced Transformer with Rotary Position Embedding, November 2023. URL http://arxiv.org/abs/2104.09864. arXiv:2104.09864 cs].

58 Shikha Surana, Nathan Grinsztajn, Timothy Atkinson, Paul Duckworth, and Thomas D Barrett. Over-confident oracles: Limitations of in silico sequence design benchmarking. In ICML 2024 AI for Science Workshop, 2024. URL https://openreview.net/forum?id=fPBCnJKXUb.

59 Felix Teufel, José Juan Almagro Armenteros, Alexander Rosenberg Johansen, Magnús Halldór Gíslason, Silas Irby Pihl, Konstantinos D. Tsirigos, Ole Winther, Søren Brunak, Gunnar von Heijne, and Henrik Nielsen. SignalP 6.0 predicts all five types of signal peptides using protein language models. Nature Biotechnology, 40(7):1023–1025, July 2022. ISSN 1546-1696. doi: 10.1038/s41587-021-01156-3. URL https://www.nature.com/articles/s41587-021-01156-3.

60 Sanjana Tule, Gabriel Foley, and Mikael Bodén. Do protein language models learn phylogeny? Briefings in Bioinformatics, 26(1):bbaf047, February 2025. ISSN 1467-5463. doi: 10.1093/bib/bbaf047. URL https://pmc.ncbi.nlm.nih.gov/articles/PMC11847157/.

61. 61 Lucrezia Valeriani, Francesca Cuturello, Alessio Ansuini, and Alberto Cazzaniga. The geometry of hidden representations of protein language models, October 2022. URL http://biorxiv.org/lookup/doi/10.1101/2022.10.24.513504.

62 Jesse Vig, Ali Madani, Lav R. Varshney, Caiming Xiong, Richard Socher, and Nazneen Fatema Rajani. BERTology Meets Biology: Interpreting Attention in Protein Language Models, March 2021. URL http://arxiv.org/abs/2006.15222. arXiv:2006.15222 cs].

63 Ria Vinod, Ava P. Amini, Lorin Crawford, and Kevin K. Yang. Trainable subnetworks reveal insights into structure knowledge organization in protein language models, June 2025. URL https://www.biorxiv.org/content/10.1101/2025.05.29.656902v1.

64 Lukas M. Weber, Wouter Saelens, Robrecht Cannoodt, Charlotte Soneson, Alexander Hapfelmeier, Paul P. Gardner, Anne-Laure Boulesteix, Yvan Saeys, and Mark D. Robinson. Essential guidelines for computational method benchmarking. Genome Biology, 20:125, June 2019. ISSN 1474-7596. doi: 10.1186/s13059-019-1738-8. URL https://pmc.ncbi.nlm.nih.gov/articles/PMC6584985/.

65 Daniel R Wong, Abby S Hill, and Rob Moccia. Simple controls exceed best deep learning algorithms and reveal foundation model effectiveness for predicting genetic perturbations. Bioinformatics, 41(6):btaf317, June 2025. ISSN 1367-4811. doi: 10.1093/bioinformatics/btaf317. URL 10.1093/bioinformatics/btaf317.

66 Banghao Wu, Bozitao Zhong, Lirong Zheng, Runye Huang, Shifeng Jiang, Mingchen Li, Liang Hong, and Pan Tan. Harnessing protein language model for structure-based discovery of highly efficient and robust PET hydrolases. Nature Communications, 16(1):6211, July 2025. ISSN 2041-1723. doi: 10.1038/s41467-025-61599-z. URL https://www.nature.com/articles/s41467-025-61599-z.

67 Wayland Yeung, Zhongliang Zhou, Sheng Li, and Natarajan Kannan. Alignment-free estimation of sequence conservation for identifying functional sites using protein sequence embeddings. Briefings in Bioinformatics, 24(1):bbac599, January 2023. ISSN 1477-4054. doi: 10.1093/bib/bbac599. URL 10.1093/bib/bbac599.

68 Zhidian Zhang, Hannah K. Wayment-Steele, Garyk Brixi, Haobo Wang, Dorothee Kern, and Sergey Ovchinnikov. Protein language models learn evolutionary statistics of interacting sequence motifs. Proceedings of the National Academy of Sciences, 121(45):e2406285121, November 2024. doi: 10.1073/pnas.2406285121. URL https://www.pnas.org/doi/10.1073/pnas.2406285121.

